# The Hippo kinase LATS1 controls CTCF chromatin occupancy and the hormonal response of three-dimensionally grown breast cancer cells

**DOI:** 10.1101/2023.11.20.566232

**Authors:** Julieta Ramirez, Roberto Ferrari, Rosario T. Sanz, Marta Valverde-Santiago, A. Silvina Nacht, David Castillo, Francois Le Dily, Maria Victoria Neguembor, Marco Malatesta, Marc A. Marti-Renom, Miguel Beato, Guillermo P. Vicent

**Affiliations:** Center for Genomic Regulation (CRG), Barcelona Institute for Science and Technology (BIST) Barcelona, Spain; Universitat Pompeu Fabra (UPF), Barcelona, Spain; CNAG-CRG, Centre for Genomic Regulation, The Barcelona Institute of Science and Technology, Baldiri Reixac 4, Barcelona 08028, Spain; Molecular Biology Institute of Barcelona, Consejo Superior de Investigaciones Científicas (IBMB-CSIC), C/ Baldiri Reixac, 4-8, 08028, Barcelona, Spain; ICREA, Barcelona, Spain

## Abstract

The cancer epigenome has been studied in cells cultured in 2D monolayers on plastic surfaces, but recent studies highlight the impact of the extracellular matrix (ECM) and the 3D environment on multiple cellular functions. Here, we report the physical, biochemical and genomic differences between T47D breast cancer cells cultured in 2D monolayer and as 3D spheroids in Matrigel. Cells within 3D spheroids exhibit a rounder nucleus with less accessible, more compacted chromatin, and altered expression of over 2,000 genes, the majority of which become repressed. Hi-C analysis reveals that cells grown in 3D exhibit enrichment in regions belonging to the B compartment, decrease on chromatin bound CTCF and increased fusion of Topologically Associating Domains (TADs). Upregulation of the Hippo pathway in 3D spheroids results in the activation of the LATS1 kinase, which promotes phosphorylation and displacement of CTCF from DNA, likely responsible for the observed TAD fusions. Cells grown in 3D exhibit higher progesterone receptor (PR) binding to chromatin, leading to an increase in the number hormone-regulated genes. This effect is in part mediated by the LATS1 activation, which favors cytoplasmic retention of YAP and CTCF removal.

## Introduction

Over the years, two-dimensional (2D) cultures have been used to improve our understanding of cellular signaling pathways and to decode the mechanisms deregulated in many diseases, including cancer. Yet, in recent years increasing evidence showed that 2D cultures do not fully reflect the complexity of the microenvironment encountered by cells as part of tissues or organs, endorsing the use of three-dimensional (3D) culture conditions generated by the presence of more physiological extracellular milieu (Kim et al., 2020). Within tissues, cells are exposed to a complex environment, including blood circulating molecules, neighboring cells, and the extracellular matrix (ECM). 3D cell culture systems have been experimented for decades (Barcellos-Hoff et al., 1989) and the importance of the ECM in cell behavior and gene expression is now well accepted.

The Hippo pathway is one of the several routes that have been reported to respond to the cell’s environment. In mammals, the core Hippo pathway is characterized by Serine/Threonine kinases; mammalian Sterile 20-related 1 and 2 kinases (MST1 and MST2) and Large Tumor Suppressor 1 and 2 kinases (LATS1 and LATS2). The transcriptional co-activators Yes-associated protein (YAP) and the Transcriptional coactivator with PDZ-binding motif (TAZ) act as nuclear relays of mechanical signals exerted by ECM rigidity and cell shape (Dupont et al., 2011). It has been previously described that on soft substrates, YAP is inactivated by LATS-dependent phosphorylation on Serine 127 and later tagged for degradation in the cytoplasm (Meng et al., 2016). The unphosphorylated YAP translocates into the cell nucleus followed by regulation of gene transcription (Zanconato et al., 2016). LATS1 and LATS2 (LATS) have emerged as central regulators of cell fate, and modulate the functions of numerous oncogenic or tumor suppressive effectors, including YAP/TAZ, the Aurora mitotic kinase family and the tumor suppressive transcription factor p53 (Furth and Aylon, 2017). Except for YAP, not many substrates have been described for LATS, however it was recently reported that LATS can phosphorylate the architectural protein CTCF within its zinc finger (ZF) linkers, reducing their affinity for DNA (Luo et al., 2020).

In breast cancer cells, steroid hormones are key regulators of cell differentiation and proliferation. The steroid hormones estrogens and progesterone acting via their specific receptors (ERα and PR, respectively) control the proliferation of breast cancer cells in a very different way. While estrogens are inducers of cell proliferation that activate cyclin D1 (Musgrove et al., 1994; Planas-Silva et al., 1999) and decrease the expression of CDK inhibitors (Prall et al., 1997), progestins promote a single cell cycle followed by proliferation arrest at G1/S, that correlates with a delayed activation of CDK inhibitor p21^WAF1^ (Owen et al., 1998).

The activated receptors of estrogens and progesterone regulate gene expression mainly by interacting with specific DNA sequences in chromatin and recruiting chromatin remodeling complexes and transcriptional coregulators (Beato et al., 1995). In addition, a minor fraction of ERα and PR is attached to the cell membrane via palmitoylation of a conserved cysteine. Binding of the hormone to this membrane-attached receptors can activate the Src/Erk/Msk1 and CyclinA/CDK2 signaling cascades (Migliaccio et al., 1996; Vicent et al., 2006; Vicent et al., 2011), leading to PR activation via phosphorylation at S294, binding to progesterone responsive elements, and modulation of chromatin structure as prerequisite for gene regulation (Beato and Vicent, 2011). One limitation of these results is that they were obtained from breast cancer cells cultured as monolayer ignoring other physiological signaling pathways, such as the Hippo pathway. Although progress has been made toward understanding the roles of LATS in tumorigenesis, the kinase substrates or downstream target proteins mediating LATS function remain largely unknown.

In the present work, we try to overcome this limitation by cultivating T47D breast cancer cells as spheroids in Matrigel (3D) and compared the results with those obtained with the same cells grown in monolayer on plastic (2D). We found that compared to 2D grown cells, the nucleus of the 3D cells is more rounded and exhibits a larger surface, a more compact chromatin and a distinctive set of 1,700 down-regulated genes. Using Hi-C assays, we did not detect strong differences at the genome structural level between cells grown in 3D and those gown in 2D, but only a modest increase in the B compartment regions and a tendency to TAD fusions. However, we did observe an increase in LATS1 kinase activity in 3D grown cells, which triggered the displacement of the architectural protein CTCF from 75% of its genomic binding regions, including the promoters of genes regulated in 3D. Despite less accessible chromatin in 3D, hormone regulation was more pronounced compared to 2D, with more responsive genes and more extensive PR binding.

## Results

### Characterization of T47D-derived spheroids

To explore how the extracellular matrix (ECM) and cell-cell contacts impact on the structure and function of the cell nucleus, we grew breast tubular epithelial cells on Matrigel, to enable the formation of three-dimensional (3D) structures. We cultured breast cancer cell lines MCF10A, T47D, MCF-7; BT-474 and ZR-75-1 on plastic (2D) or embedded on Matrigel (3D), for 10 days, the time required to form multicellular acini (Appendix Figure S1A). While T47D cells formed spheroids containing proliferating cells, the non-tumorigenic MCF10A cells formed round structures with a polarized acinar internal pattern (Figure 1A). The other tumorigenic cell lines (MCF-7 and BT-474) cultured on Matrigel also form spheroids as previously reported (Lee et al., 2007) (Appendix Figure S1A). In this study, we focused on PR+ T47D cells as a model system. These cells are known for their robust responsiveness to the hormone progesterone, and their genomic structure, the PR cistrome and the associated signaling pathway network have been extensively characterized (Ballare et al., 2013; Le Dily et al., 2014; Le Dily et al., 2019; Vicent et al., 2006; Vicent et al., 2010; Zaurin et al., 2021).

**Figure 1.**
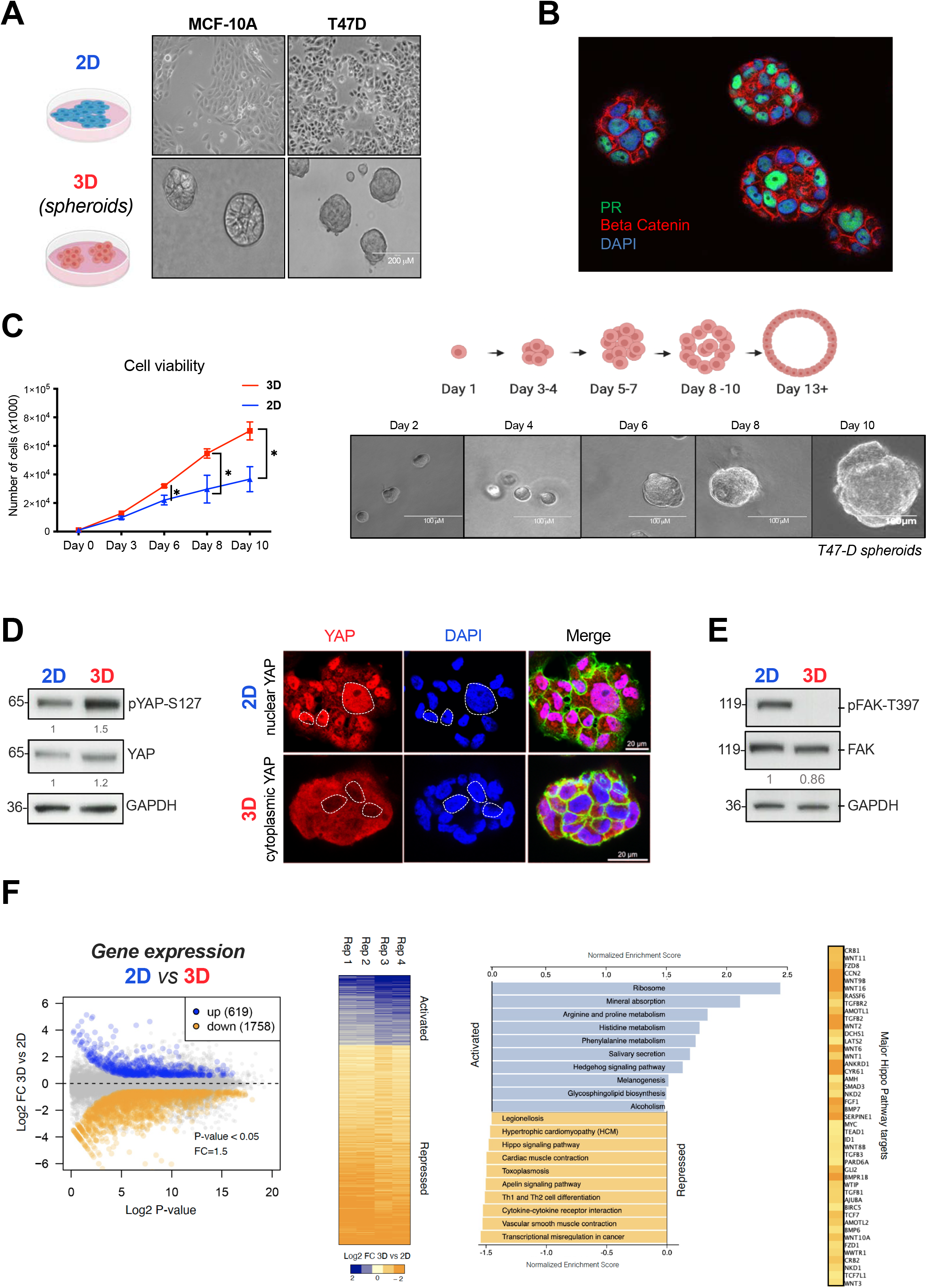
Evaluation of cell growth, signaling pathways and differentially expressed genes in T47D cells grown in 3D and 2D conditions. **A.** Comparison of MCF10-A and T47D breast cell lines grown on the plastic dish (2D) and as 3D culture in Matrigel. **B.** Immunofluorescence analysis showed that PR expression is conserved in 3D T47D cells. **C.** Cell proliferation assays performed in 2D and 3D cells (left panel). The growth of 3D T47D cells over a period of 10 days to reach an estimated size of 100 µm is depicted (right panel). **D.** Increased levels of YAP protein phosphorylation (pYAP-S127) in 3D was assessed by western blot (left panel). Immunostaining of Hippo pathway effector YAP in 2D and 3D T47D cells. When cells are grown in 3D conditions YAP is phosphorylated and re-localized to the cytoplasm (right panel). **E.** T47D cells in grown in a 3D down-regulate phosphorylation of FAK and its downstream pathways. **F.** RNA-seq Differential Expression Analysis (DEA), performed in 2D and 3D T47D cells using [log2FC=2 and p value<0.05], the experiment reveals over 2,300 genes deregulated when cells were grown in 3D (left panel). Using the DEA results 3D/2D, enriched signaling pathways (KEGG) are shown, FDR <0.05 (right panel).

We performed a functional and phenotypic characterization of the T47D spheroids. First, we assessed the physical properties of the cell nuclei in the spheroids compared to 2D by implementing IMARIS and ImageJ tools (Martin et al., 2021). The nuclear volume of 3D cells, measured after DAPI staining, was significantly enlarged compared to the 2D cell nuclei (488.7 μm^3^ vs. 408.5 μm^3^, p value<0.001, Appendix Figure S1B, left panel) while the diameter of the 3D nuclei was smaller (9.17 μm vs.15.56 μm, p value<0.001, Appendix Figure S1B, middle panel). This translates into an increased sphericity coefficient of the 3D nuclei (0.74 vs. 0.57 in 2D, p value<0.001, Appendix Figure S1B, right panel) and a larger surface area (643.3 μm^2^ vs. 381.6 μm2, p value<0.001, Appendix Figure S1B, lower panel).

We also evaluated the proliferation capacity of T47D cells cultured in 3D compared to cells grown in 2D. We found that 3D cells proliferated at a rate 2.6 times faster than its 2D counterpart (Figure 1C, left panel). Within 10 days of culture the 3D cells formed spheroids with an average diameter of 100 μm, typically comprising 119± 12 cells (Figure 1C, right panel). The increased cell proliferation detected in 3D cells was in accordance to the high levels of Ki67 positive cells detected in the spheres, also evident by using a live/dead cell imaging Kit (Appendix Figure S2A).

Next, we probed the signaling pathways known to be regulated in cells grown in 3D conditions, such as the Hippo and the Focal Adhesion Kinase (FAK/ PTK2) pathways. In 3D cultured breast cancer cells, YAP displayed higher level of phosphorylation on S127 compared to 2D, while total YAP levels remained unchanged (Figure 1D, left panel). YAP was also found largely in the cytoplasm of 3D grown cells (Figure 1D, right panels), as previously described (Zanconato et al., 2019). In contrast, in 2D cells, the Hippo pathway was switched off and YAP mainly found into the nucleus (Figure 1D, top right panels). When probing the FAK/PTK2 pathways, we found p-FAK severely deregulated in 3D cells, as indicated by its significant reduction in FAK-Y397 phosphorylation, compared to the levels found in 2D cells (Figure 1E), despite no changes in total FAK levels (Figure 1E). Therefore, in 3D grown cells the presence of a less stiff environment is sensed, integrated and transduced leading to both repression of focal adhesion and activation of the Hippo pathway.

Increased levels of phospho-LATS (part of the Hippo pathway) in response to 3D cell growth was also observed in ER^+^ MCF-7 cells. However, this activation was less pronounced in comparison to T47D cells (Appendix Figure S1C compared to Figure 5B). Notably, there was no significant increase in p-YAP or decrease in p-FAK in MCF-7 cells, where YAP expression notably remained low in comparison to other mammary cell lines (Appendix Figure S1C, lower panel). This observation indicates a partial conservation of the cytosolic activation of the Hippo pathway in response to 3D in MCF-7 cells. (Appendix Figure S1C). Additionally, it is worth mentioning that reduced ER levels in MCF-7 3D cells (new Appendix Figure S1C) had an impact on their proliferative capacity (data not shown).

To investigate whether these changes in signaling, shape, surface and volume of the 3D nucleus impact on gene activity, we performed RNA-seq analysis in T47D cells grown in 2D and 3D. The transcriptional profiles of 2D and 3D samples, yielded statistically significant changes for 2,377 transcripts: 619 were up-regulated and 1,758 down-regulated in 3D cultured cells compared to 2D cells [log2 FC=2 and p value<0.05] (Figure 1F). Gene Ontology analysis of biological processes and cellular compartments showed that cultivating cells in 3D lead to up-regulation of genes enriched in terms associated to neurogenesis, nervous system development, axon genesis and neuronal projection. While 3D down-regulated genes were overrepresented in cell and focal adhesion, cell junction and cell periphery (Figure EV1A). These results suggest that cells cultured in 3D possess a unique gene expression program that recapitulates neuronal growth and, additionally, decreases the expression of genes associated to adhesion, most likely due to the presence of a less stiff environment compared to 2D. The similarities between growth as 3D spheres and the nervous system has been previously proposed (Caswell and Zech, 2018). Further analysis of KEGG exclusive 3D pathways showed terms related to *Ribosome*, *Mineral Absorption* and *Amino acid Metabolism* for 3D up-genes (Figure 1F, right panel). The physical characteristics of the surrounding structure play a pivotal role in coordinating this regulatory interplay. Concerning *Amino acid Metabolism*, the stiffness of the extracellular matrix regulates the activation of the Hippo pathway and, consequently, the nuclear localization of YAP/TAZ. YAP/TAZ play a critical role in controlling genes involved in amino acid synthesis, transport and metabolism (Ge et al., 2021). Therefore, in response to the mechanical properties of their environment, cells adjust amino acid metabolism to acquire the energy necessary for specific cellular functions, such as enhanced cell proliferation, as illustrated in Figure 1C.

Extracellular matrix stiffness also influences *Mineral Absorption*, as previously documented (Derricks et al., 2015). Cells on softer substrates (4 kPa) demonstrate increased responsiveness to VEGF, while cells on stiffer substrates (125 kPa) exhibit a diminished response.

The enrichment of genes related to *Ribosomes* suggests that protein synthesis is differentially regulated in 3D environments. This may be attributed to the unique demand for specialized proteins involved in cell-cell interactions, the integration of external signals, and mechano-transduction, especially in cells growing as spheres with low stiffness. In summary, the regulation of genes associated with *Amino acid Metabolism*, *Mineral Absorption*, and *Ribosomes* in 3D environments underscores the significant influence of extracellular matrix stiffness on various cellular processes. This intricate interplay allows cells to adapt to their mechanical surroundings, ultimately affecting crucial aspects of cell biology and physiology.

The 3D-repressed genes exhibit an enrichment in genes associated with *Transcriptional deregulation in Cancer*, the *Hippo* and *Apelin pathways*, among others. The Apelin pathway is not well-documented in breast cancer, with only one report associating high levels of Apelin with postmenopausal breast cancer (Salman et al., 2016). However, genes involved in vasodilation and *Muscle Contraction* are significantly repressed, which could be explained as an adaptation to the softer 3D environment. The suppression of these pathways could also be linked to the up-regulation of *Mineral Absorption* pathway, as calcium ions (Ca2+) can regulate the contractility of vascular smooth muscle cells (Brozovich et al., 2016).

Next, we asked whether the changes in gene expression that we observed in 3D grown cells (Figure 1F) were due to changes in histone acetylation. To this end, cells grown in 2D and 3D were subjected to ChIP-seq of H3K27ac and H3K18ac, two marks strongly associated to gene expression (Ferrari et al., 2012; Wang et al., 2008). In general, a notable variation, ranging from 20% to 44%, was observed in the two marks when cells transitioned from 2D to 3D culture conditions (Figure EV1A-B). These alterations were particularly concentrated in genes influenced by the growth environment. Consequently, a substantial reduction in H3K27ac was detected for genes down-regulated and up-regulated in 3D (Appendix Figure S3C), along with an increase in H3K18ac for genes up-regulated in 3D (Appendix Figure S3D, right panel), as compared to their 2D counterparts.

RNA polymerase 2 (pol2) which globally didn’t show drastic changes (Appendix Figure S3E) was significantly reduced at the TSS and gene body of 3D down-regulated genes (Appendix Figure S3F, left panel) and increased in 3D up-regulated genes (Figure S4F, right panel) as visualized at the *HSPA8* and *CXCL12* loci, two representative genes of each category (Appendix Figure S3G).

We next tested whether the levels of repressive marks such as H3K27me3 and H3K9me3 are affected on genes that change their expression with the culture condition. In 3D down-regulated genes an increase in H3K27me3 was detected (EV2A) while H3K9me3 levels remained unchanged (EV2B). Surprisingly, a significant increase in H3K9me3 was found in in 3D-up genes (EV2B).

Given that Polycomb operates through a digital mechanism with two opposing expression states signifying its presence or absence on target genes (Menon et al., 2021), assigning the modest increase in H3K27me3 detected in 3D down genes directly to its final impact on transcription is challenging. Instead, it is likely a complex interplay of various histone modifications, along with other transcriptional and architectural factors, that contributes to this phenomenon.

We also evaluated whether the change in cell culture condition affects the presence and distribution of super-enhancers (SEs) (Hnisz et al., 2013). To date, 214 genomic regions were identified as SEs in T47D breast cancer cells (Jiang et al., 2019). The overlap between the super-enhancers detected in 2D and 3D according to the H3K27ac signal (579 and 564, respectively) was close to 80% (Figure EV2C). This observation would support that cells fundamentally maintain their regulatory gene network in both of these environments (Figure EV2D). Nonetheless, it’s essential to acknowledge that 20% of super-enhancers are exclusively activated in the 3D environment. The question of whether these variances have implications for the alteration of cell identity remains open. To address this point, by using a proximity-based script (Hnisz et al., 2013), we found 129 and 95 genes associated to 2D and 3D SEs, respectively (Figure EV3A). The 2D-exclusive genes are related to terms like abnormal embryo size, abnormal development, and hematopoiesis, which are closely linked to the Hippo pathway (as reported in (Yu et al., 2015)) (Figure EV3C). Conversely, in the case of genes exclusively regulated in the 3D condition by SEs, many of them appear to be artifacts, as illustrated with the HSPA8 gene (Figure EV3B). This hampers the identification of significant associated terms.

When attempting to establish a connection between super enhancers and alterations in genome structure, such as the A to B transition, it becomes evident that the quantity of SEs is reduced, and this transformation exhibits subtler changes, predominantly shifting from A to less degree of A, or toward emerging B, and vice versa (Figure EV3D). Snapshots from the genome browser illustrating these transitions for two 3D down-regulated genes, SLC26A7 and RUNX1T1, and two 3D up-regulated genes, PI15 and ITGAL are shown (Figure EV3D).

### Impact of the 3D growth on nuclear structure

To explore how 3D culture impacts on nuclear structure, we measured the distribution of the chromatin throughout the nucleus by determining the coefficient of variation of DNA stained with DAPI (Martin et al., 2021). We found that the chromatin is organized differently, presenting a more heterogenous pattern in 3D grown cells, with an increased coefficient of variation compared to cells grown in 2D (Figure 2A, left panel), a broader distribution of DAPI signal intensity and the appearance of highly dense foci in 3D grown cells (Figure 2A, middle and right panels, respectively).

**Figure 2.**
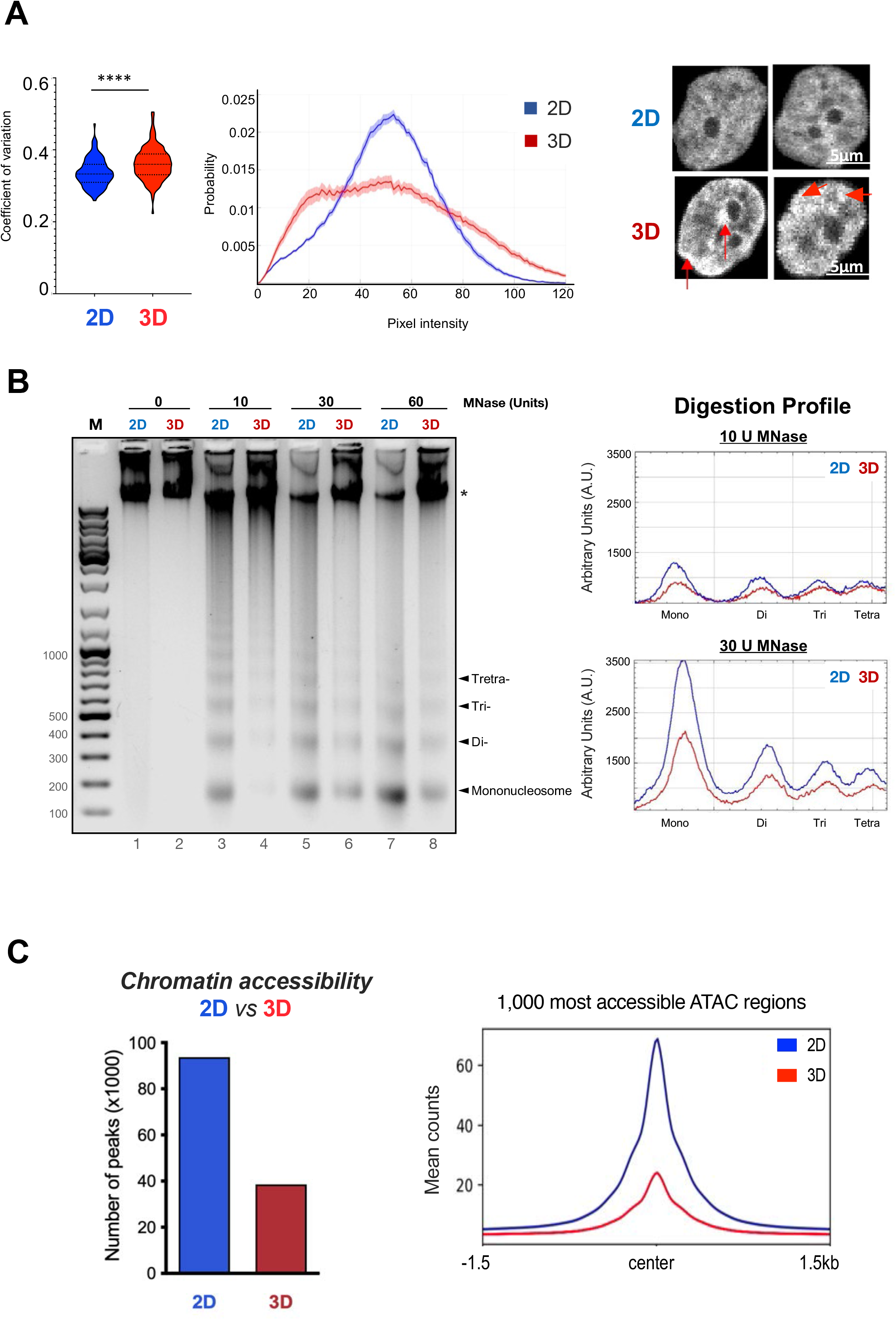
Chromatin organization and accessibility in 2D and 3D grown cells. **A.** Coefficient of variation of DAPI staining distribution between T47D cells grown in 2D and in 3D, n= 100 cells per condition. Violin plot (left panel). Probability distribution of the pixel intensities obtained in 2D and 3D (middle panel). T47D nuclei stained with DAPI exhibiting a clearer pattern of condensed chromatin clusters in 3D grown cells is shown (right panel, red arrows indicate highly compacted regions). Scale bar: 5 μm. **B.** T47D cells grown in 2D and 3D conditions were isolated, nuclei were prepared and digested with different concentrations of MNase. Digestion products were resolved in 1.2 % agarose gel electrophoresis (left panel) and the intensity of the bands was quantified by ImageJ software (right panel). **C.** Number of peaks per sample of ATAC-seq experiments (up left panel). The 1,000 peaks with the highest score for 2D and 3D condition were plotted (bottom left panel). The overlap between ATAC-seq replicates is shown (right panel).

In order to assess chromatin accessibility within the cell nucleus of 2D and 3D-grown cells, we conducted a comprehensive approach that includes: i) MNase experiments, which detect the compaction of chromatin fibers at a large scale, potentially indicating changes in the presence of linker histones and heterochromatin, and ii) ATAC-seq experiments, which measure focal nucleosome depletion at sites where sequence-specific transcription factors, such as CTCF, bind, as well as the nucleosome-depleted region at transcription start sites (TSS).

To estimate the accessibility of the chromatin at large scale, we performed micrococcal nuclease (MNase) digestion assays. Isolated nuclei obtained from cells grown in 2D and 3D were treated with increasing concentrations of MNase, and the products of digestion were analyzed by gel electrophoresis (Figure 2B). We found that the accessibility to MNase was compromised in 3D compared to its 2D counterpart at the range of MNase concentrations tested. (Figure 2B). In addition, the amount of undigested DNA present in 3D was higher compared to 2D, even at 60U MNase (Figure 2B, left panel). These results support that chromatin is less accessible in nuclei from 3D growing cells.

To assess whether the chromatin accessibility changed at nucleosome level with the growth conditions, we performed an Assay for Transposase-accessible chromatin followed by sequencing (ATAC-seq), which is a technique used to assess genome-wide chromatin accessibility (Buenrostro et al., 2013).

Nuclei obtained from cells grown in 2D and 3D conditions were incubated with 2.5 U of the tagmentase Tn5. The average number of ATAC peaks was significantly higher in 2D than in 3D growing cells (93,850 vs. 38,542 peaks for 2D and 3D, respectively) (Figure 2C), pointing to a general decrease of accessibility in nuclei from 3D grown cells. When comparing the 1,000 most accessible ATAC regions detected in both 2D and 3D grown cells, we observed that 3D peaks contained less reads/peak (Figure 2C, right panel). Most of the peaks detected in 3D grown cells also appeared in 2D (96.88 %; 37,342). Only 547 new ATAC peaks were detected in 3D grown cells, while 24,242 sites were lost when comparing all replicates. Hence, the chromatin structure in 3D-grown T47D cells exhibited a noticeably less accessible conformation, both at the level of larger fibers and at nucleosome resolution, in contrast to the 2D cells. These findings align with the results from gene expression analysis and imaging studies (Figure 1F and Figure 2A-C).

To characterize the regions that become more accessible in 2D and 3D we conducted a more in-depth analysis of the ATAC-seq data (EV4A-B). Our results suggest that the sites that become more accessible in 2D and 3D are enriched in the TF motifs of CTCF, TEAD, FOXA1 and PR motifs (EV4B). However, when we combined our analysis with ChIP-seq data (Zheng et al., 2019), word cloud plots indicated that open sites in 3D are enriched in the estrogen pathway through the ER itself, alongside pioneer factors FOXA1 and GATA3; the cofactor GREB1 and the coactivator P300 (EV4B-C, lower panels), which could explain the increased proliferation observed in 3D cells (Figure 1C). In contrast, the sites that become closed in 3D (2D-exclusive) are enriched in CTCF, an architectural factor that we have shown to be displaced when we grow the cells as spheroids (Figure 4A). This would imply that in 3D cells a more estrogenic program is turned on along CTCF is displaced, if these two events are connected require further investigation.

Furthermore, the chromatin accessibility of 2D and 3D-exclusive genes was also assayed, and we detected changes. As expected, 2D-up genes exhibit greater accessibility within the gene body in 2D condition compared to their 3D counterpart, whereas in 3D condition the expected inverse trend is not observed (EV4D). However, we found increased accessibility in distantly located regulatory regions (enhancers) associated to 166 genes up-regulated in 3D. To illustrate this finding, the TN2Cx and PTPRD genes are shown (EV4E).

### Effect of the 3D growth on genome architecture

Next, we addressed the overall 3D genome organization performing Hi-C experiments with cells grown in 2D and 3D conditions.

First, we explored the chromosome compartments by analyzing the correlation between the eigen vectors obtained from the interaction of Hi-C matrices in 2D and 3D cultured cells at 1 Mb resolution. This analysis showed no substantial changes in the genome structure at this resolution (R=0.99) (data not shown). We next evaluated the genome compartmentalization at 100 kb resolution and found a limited number of changes between cells grown in 2D and in 3D. We identified 50 regions that changed from A (active) to B (inactive) compartment in 3D grown cells, while only nine regions changed from B to A compartment (Figure 3A). In fact, when the A/B compartment transitions were evaluated by calculating an A/B score using Hi-C homer (Heinz et al., 2018), we observed significant increase of 3D up genes in A compartments and conversely, decrease in A and increase in B compartments for down-regulated genes in 3D (Figure 3B). These findings suggest that in 3D condition the changes detected in gene expression sustain gradual transitions between the A and B compartments.

**Figure 3.**
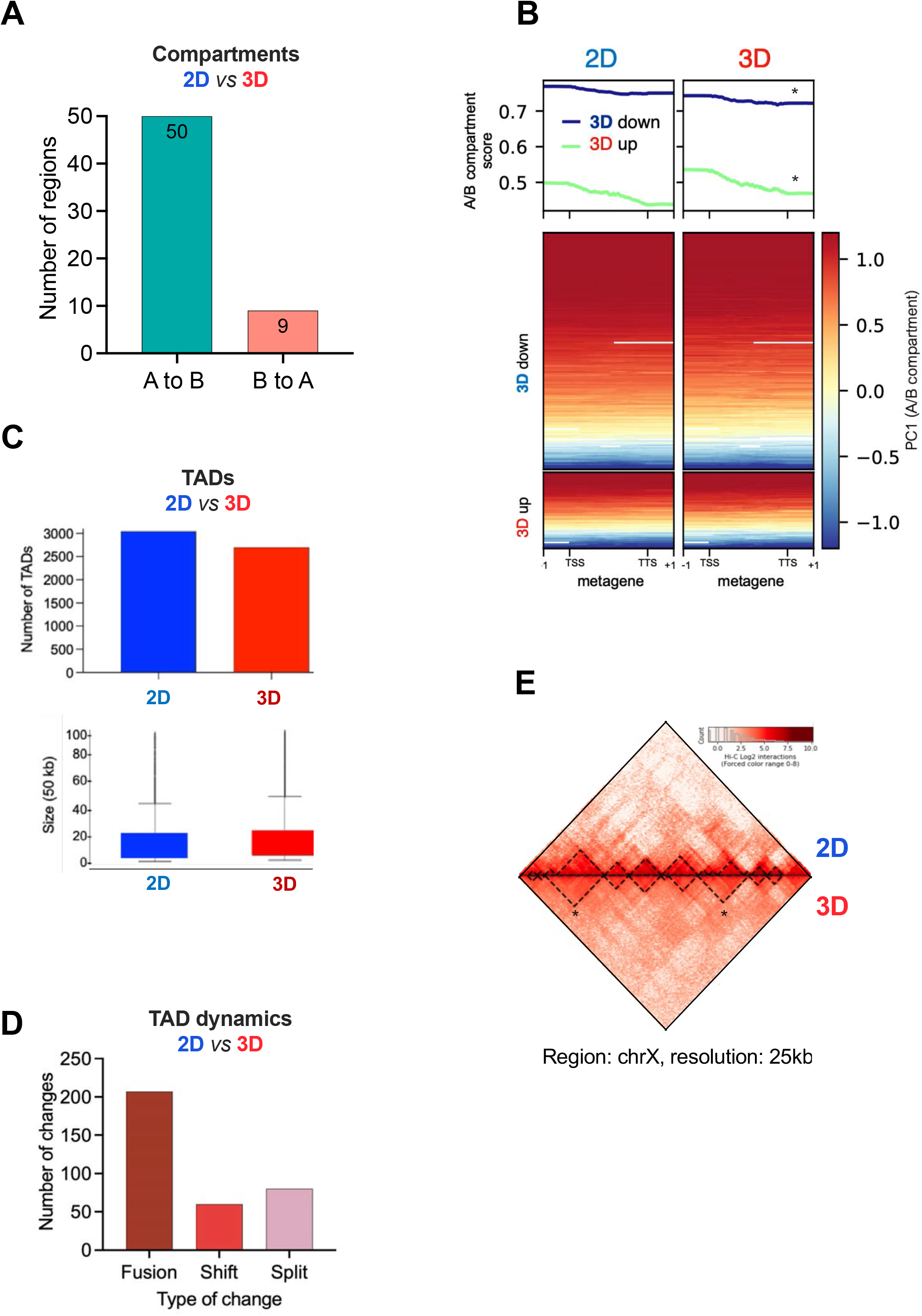
Genome topology in 2D and 3D-grown cells. **A.** Total number of compartments changing from 2D to 3D cells (bin= 50 kb) **B.** Metagene analysis of A/B compartments versus gene expression. Metagene profiling of A/B compartments for 3D down and 3D up genes is shown both as average profile (top) and heatmap (bottom). Both sets of genes show significant changes in compartment score (* tow tail t-test p-values < 0.001). **C.** Changes in total number (top) and size of TADs (bottom) between 2D and 3D grown cells. **D.** Type of TAD changes detected in 3D grown cells. **E.** A Hi-C heatmap of a region located in the X chromosome illustrating TADs fusions in 3D.

Finally, we assessed the behavior of topologically associating domains (TADs). As depicted in Figure 3C, in 3D grown cells there is a decrease in total number of TADs (3,041 and 2,860 in 2D and 3D, respectively) accompanied by a small increase in their size. This is due to the prevalence of TAD fusions (200) over splitting (60) or shifting (50) (Figure 3D). A Hi-C heatmap plot at chromosome X depicting two TAD fusions detected in 3D grown cells is shown (Figure 3E).

Since the discovery of TADs, it became clear that the boundary regions separating topological domains are enriched in CTCF along with the structural maintenance of chromosomes (SMC) cohesin complex, housekeeping genes and SINE elements (Barutcu et al., 2018; Dixon et al., 2012; Nora et al., 2012).

CTCF is a highly conserved zinc finger protein and is best known as a transcription factor. It can function as a transcriptional activator, repressor or as an insulator protein, blocking the communication between enhancers and promoters (Kim and Wirtz, 2015). Thus, targeted degradation of CTCF can affect either enhancer-promoter looping or local insulation, promoting TAD fusions. To map the genome distribution of CTCF in both 2D and 3D conditions, we performed ChIP-seq experiments. Even though CTCF protein levels remain unchanged in 2D and 3D growing conditions (Figure 4A, bottom right panel), we found a 75 % loss of CTCF binding in 3D cells (Figure 4A, top right panel). To rule out that the antibody against CTCF might not be properly recognizing CTCF in 3D, we perform chromatin fractionation assays in 2D and 3D grown cells. Our results showed that about half of the CTCF is bound to 3D chromatin fraction compared to the 2D condition (Appendix Figure S4A).

**Figure 4.**
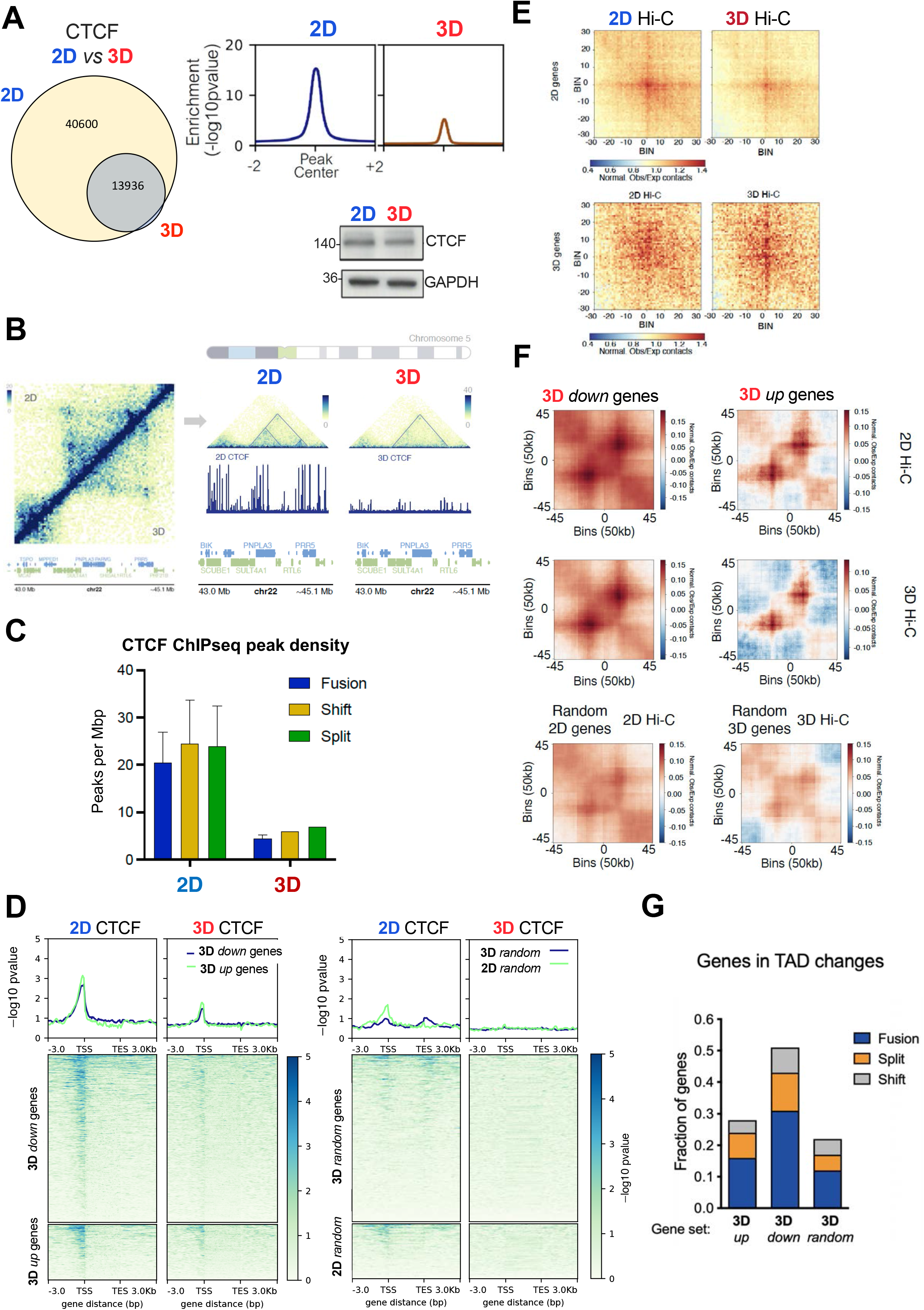
The architectural protein CTCF is displaced from target chromatin in 3D grown cells. **A.** Cells grown in 2D and 3D were lysed and total extracts were analyzed by western blot using CTCF and GAPDH-specific antibodies (left panel). ChIP-seq of CTCF performed in 2D and 3D conditions showed differential CTCF binding regions genome wide (middle and right panels). **B.** Snapshot of a region presenting a 3D-exclusive TAD fusion event that overlaps with the loss of CTCF binding. **C.** The density of CTCF peaks in 3D-exclusive fused, split and shifted TADs is shown. **D.** The presence of CTCF in 3D up- and down-genes (left panels) and in random genes (right panels) are shown. **E**. Hi-C explorer aggregate plots. Long-distance interactions among 3D down- and up-regulated genes grown in 3D and 2D conditions. The genomic coordinates of the 3D down and up-regulated genes are centered between half the number of bins and the other half number of bins. Plotted are the submatrices of the aggregated contact frequency for 20 bins (1.5 kb bin size, 35 kb in total) in both upstream and downstream directions. Color bar scale with increasing red shades of color stands for higher contact frequency. **F.** Aggregate FAN-C plots depict a region that is three times the size of the TAD located in its center. TADs are selected on the basis of containing 3D up-or down-genes (upper panels) or random genes (lower panels). High signal is located in the center, especially at the TAD corner, where the corner loops are typically located. **G.** CTCF displacement impacts in 3D-regulated genes, by changing their contact environment through TADs fusion. The percentages of up, down or random genes in 3D condition that fall into fused, split or shifted TADs are shown.

This led us to hypothesize that the loss of CTCF binding could be responsible for the loss of TAD borders resulting in their fusion detected in 3D grown cells (Figure 3E). We found regions where TAD fusions and CTCF displacement overlap in 3D (Figure 4B). However, the loss of CTCF binding was global and distributed throughout the entire genome, impacting on all TADs, irrespective of whether they change or not in 3D (Appendix Figure S4B-D). Similar loss of CTCF was found at fused, shifted or split TADs (Figure 4C). In fact, we identified several regions where the loss of CTCF did not have impact on the TAD structure at all (Appendix Figure S4E). Around 95% of the TAD borders were conserved in 2D and 3D conditions.

To assess whether changes in the 3D TADs could be assigned to differential loading of cohesin components in these regions, we conducted RAD21 ChIP-seq experiments under 2D and 3D growth conditions. Our findings indicate that there are no significant alterations in RAD21 occupancy within both fused and random TADs (Appendix Figure S5).

Interestingly, HOMER motif analysis of the ATAC-seq peaks showed that the motif for CTCF tops the rank of enrichment in sites where accessibility is reduced in 3D (Appendix Figure S4F). It is important to note that even though BORIS (CTCFL) is ranked as second, this protein is not present in breast cancer cells. Several reports show that displacement of CTCF leads to a decrease in ATAC-seq signal and gene regulation (Franke et al., 2021; Xu et al., 2021). This would suggest that in 3D, CTCF displacement results in a more compact chromatin at those sites and could affect gene activity.

### CTCF displacement and gene regulation in T47D 3D cells

Although the displacement of CTCF observed in the chromatin of 3D-grown T47D cells does not always lead to changes in TADs boundaries examined by Hi-C, we wanted to investigate whether the absence of CTCF is associated to changes in gene regulation. Combined analyses of RNA-seq and ChIP-seq data showed that 3D up and down-regulated genes showed a significant depletion in CTCF around the TSS compared to random genes (Figure 4D). Therefore, the genes activated or repressed as a result of growing cells in 3D, present less CTCF around the transcription start site, which would imply a reduction of long-distance interactions (“looping”) between regulatory regions (enhancers, silencers) and their target genes. In fact, this loss in interactions detected in 3D resulted into a decrease of Hi-C contacts, particularly evident in 3D down-regulated genes (p-val < 0.001) (Figure 4E and S6G).

As mentioned above, CTCF is also enriched at TADs borders. We asked whether long range CTCF-interactions and therefore TADs structure as a whole, could be particularly affecting those TADs specifically enriched in 3D de-regulated genes. To address this point, we used FAN-C aggregate plots using the TADs containing genes differentially expressed in 3D grown cells and measure the levels of CTCF by ChIP-seq. Our results showed that for genes down-regulated in 3D, the global interactions between two CTCF sites that formed a TAD corner were drastically reduced (p-val < 0.001) (Figure 4F, left panels), whereas for genes activated in 3D cells we could detect a slight decrease, but not statistically significant (p-val > 0.001) (Figure 4F, right panels). No difference was found for a random set of genes and TADs as a control (Figure 4F, bottom panels). Therefore, CTCF displacement could be involved in the prevailing 3D gene repression. This implies changes at the level of intra-TAD interactions as well as at the level of TAD borders, decreasing its ability to isolate and regulate genes.

We then assessed whether the observed differences in CTCF and TAD structure associated to 3D genes (Figures 4E-F) could lead to any preferential change of TADs in 3D condition. We found that 17% of up and 30% of down-genes in 3D are included in 3D fused TADs, while the percentages were significantly lower in split and shift TADs (7 and 13% for split and 3.5 and 7% for shifted TADs, P: 0.001) as well as in all categories for random genes (Figure 4G). Thus, CTCF displacement particularly impacts in 3D genes, by changing their contact environment through TADs fusion as shown for the 3D down gene *DUSP1* whose expression is associated with an increased risk of metastasis and shorter overall survival in breast cancer (Candas et al., 2014) (Appendix Figure S6A-C).

### LATS activity and CTCF binding to chromatin in T47D grown in 3D cells

In 3D condition, we have shown that the Hippo pathway is activated, resulting in the phosphorylation of YAP at serine 127 by the LATS kinase (Figure 1D).

A recent study reported that LATS can phosphorylate CTCF at T347 and S402 and disables its DNA-binding activity (Luo et al., 2020). In that system, loss of CTCF binding was able to disrupt local chromatin domains and down-regulate genes located in the neighborhood (Luo et al., 2020).Therefore, we tested whether the displacement of CTCF found in 3D grown cells could be due to increased LATS-dependent phosphorylation of CTCF.

As no commercial phos-CTCF antibody is available, we performed a Phos-tag gel, that would allow detection of phos-CTCF as slow migrating bands depending on their state of phosphorylation. When extracts from 2D and 3D grown cells were analyzed in Phos-tag gels, we detected a retarded band in 3D, which was not present in 2D extracts (Figure 5A). To confirm that this band corresponded to a phosphorylated version of CTCF, we prepared 3D extracts by using a buffer lacking phosphatase inhibitors. Under this condition, the CTCF band ran faster in Phos-tag gel as a non-phosphorylated version of CTCF (compare lanes 2 and 3 in Figure 5A), supporting that phosphorylated-CTCF is preferentially found in extracts from 3D grown cells.

**Figure 5.**
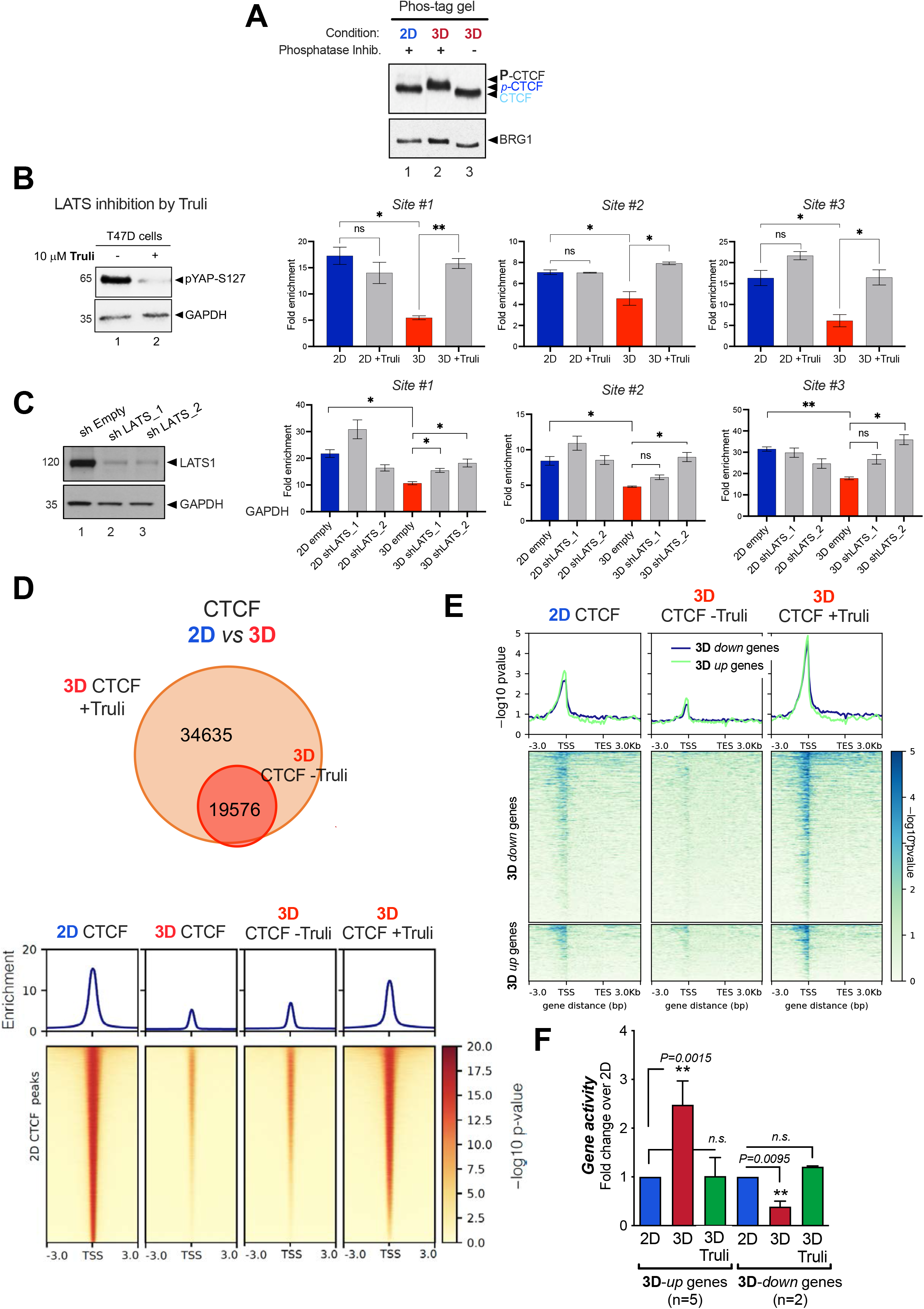
Inhibition or depletion of the Hippo pathway LATS kinase compromises the loss of CTCF detected in 3D grown cells. **A.** Cells grown in 2D and 3D were lysed and total extracts were analyzed by Phos-tag gel which identifies putative phosphorylated bands of CTCF exclusively in 3D. **B.** Cells grown in 2D or 3D conditions and treated or not with the LATS inhibitor TRULI were subjected to ChIP-qPCR of CTCF in three regions where CTCF binding is lost in cells grown in 3D (n=2, p value <0.05) (right panel). T47D cells treated or not with TRULI were lysed and total extracts were analyzed by western blot using pYAP-S127 and GAPDH-specific antibodies (left panel). **C.** Effect of LATS-1 depletion on CTCF binding. Left panel: quantification of LATS1 by western blot. Right panel: ChIP qPCR analysis of CTCF in extracts derived from control cells transfected with shEmpty and two distinct shLATS. **D.** Cells grown in 3D conditions treated or not with the LATS inhibitor TRULI were subjected to CTCF ChIP-seq. Venn diagrams showed the overlapping in CTCF binding. **E.** Cells grown in 3D conditions and treated or not with TRULI were subjected to CTCF ChIP-seq. The enrichment of CTCF in 3D up- and down-genes as well as in random genes is shown. **F.** Cells grown in 2D and 3D conditions and treated or not with TRULI as indicated, were submitted to gene activity assays. Several up and down-regulated genes in 3D were tested. Results are represented as mean and SD from three experiments performed in duplicate. The p-value was calculated using the Student’s t-test.

The direct impact of LATS on CTCF phosphorylation was tested using TRULI, a potent ATP-competitive inhibitor of LATS kinases (Kastan et al., 2021). Incubation for 24 h with 10 µM TRULI decreased the phosphorylated (activated) version of YAP at S127 (Figure 5B, left panel).

To have a formal proof that LATS1 phosphorylates CTCF in 3D cells, we performed immunoprecipitation of CTCF in 3D-grown cells treated or not with TRULI and then probed with specific antibodies recognizing the phosphorylated RXXpS/T residues that match the changes observed in T374 and S402 as previously described (Luo et al., 2020). Our results showed that the phospho-CTCF signal in 3D is reduced by 40% in the presence of TRULI, confirming the raised hypothesis (**EV5A**).

To confirm that T374 and S402 of CTCF are the phosphorylation targets of LATS, and to assess their impact on chromatin binding in our experimental system, we transfected T47D cells with both wild-type and the phospho-mimetic T374E/S402E CTCF variant. Subsequently, we evaluated their chromatin binding abilities. The T374E/S402E mutant displayed a diminished capacity to bind to chromatin when compared to the wild-type CTCF (**EV5B**). These results confirm previous studies (Luo et al., 2020) and provide more precise insights into the LATS target sites.

It has been reported that the overexpression of CTCF impedes the proliferation and metastasis of breast cancer cells by deactivating the nuclear factor-kappaB pathway in breast cancer cells (Wu et al., 2017). To elucidate the function T474 and S402, we conducted a functional assay by evaluating cell growth in T47D cells expressing both wild-type (WT) and a phospho-mimetic variant of CTCF T374E/S402E (EV5C). The overexpression of WT CTCF in T47D cells not only inhibited growth but also triggered cell death, compared to the control transfection (EV5C). Conversely, in the T374E/S402E mutant in which chromatin binding is compromised (see EV5B), this effect was not appreciated (compare WT vs. mutant in EV5C). This suggests that LATS1-dependent phosphorylation of CTCF induced its displacement and participates in 3D cell growth, as demonstrated in Figures 1C and Appendix S8A-B.

Therefore, we performed qPCR-ChIP of CTCF in 2D and 3D grown cells in the absence or in the presence of TRULI. As expected, a decrease in CTCF binding was detected in 3D compared to 2D in the three different genomic regions tested (Figure 5B, right panels). In the presence of TRULI, CTCF binding was significantly recovered (Figure 5B, right panels).

To support these findings and to discard indirect effects of TRULI, we performed similar ChIP experiments in cells deficient of LATS1 (Figure 5C, left panel). As with TRULI, knockdown of LATS1 rescues CTCF binding in 3D cells (Figure 5C, right panels). Interestingly, these results point to LATS1 as responsible for the observed effects.

The potential involvement of LATS2 was assessed using a specific antibody that recognizes phosphorylation at T1079 and T1041 present in both LATS1 and LATS2, in wild-type cells and LATS1-depleted cells. The dual phosphorylation signal in 3D cells decreased proportionally with the reduction of LATS1 (Figure 5C), supporting that LATS1 is the Hippo kinase involved in our system (**EV5D).**

To extend our conclusions we performed ChIP-seq of CTCF in 3D cells in the presence and absence of TRULI. We found that binding of CTCF was recovered across the genome in the presence of the LATS inhibitor (Figure 5D). In fact, in this condition we observed that CTCF binding was preferentially recovered at 3D-dependent genes (Figure 5E).

The sensitivity of CTCF ChIP-seq experiments is notably influenced by the particular protocol employed. This results in varying outcomes regarding the detected 3D shift. Notably, using a stringent ChIP protocol, which includes multiple washes and the use of DNA purification columns (ChIP-IT, Active Motif), revealed a 75% displacement of CTCF when cells were cultured in a 3D environment, as depicted in Figure 4A.

In contrast, using a standard protocol with gentler wash steps, as outlined by Vicent et al., 2014 (Vicent et al., 2014), and incorporating an internal Drosophila spike-in control, reduced this displacement to 40% (Appendix Figure S7A-C). Notably, this reduction remained statistically significant when compared to cells cultured in a 2D monolayer setting. Moreover, in spike-in-controlled CTCF ChIP-seq experiments conducted in the presence of the LATS inhibitor TRULI, CTCF binding was restored (refer to Appendix Figure S7C). These findings align with our earlier results; the average of all three CTCF ChIP-seq replicates obtained from 2D and 3D cells is depicted in Figure S7D.

We next asked whether the activity of the LATS kinase and its effect on CTCF binding was required for the proper regulation of 3D genes. We tested 5 genes up and 2 genes down in 3D and we found that TRULI significantly changed their activity, approaching to that detected in 2D (Figure 5F).

Moreover, we also evaluate the effect of TRULI on 3D growth. In the presence of TRULI, 3D cells grew less and the spheres were smaller compared to untreated 3D cells (Appendix Figure S8A-B). In agreement with the gene activity, cell growth in the presence of TRULI turned out to be similar to that observed in 2D, coinciding with the presence of CTCF bound to the target genes and the activity exhibited in both conditions (Figure 5E-F and Appendix Figure S8).

### Hormone-dependent gene regulation in 2D and 3D T47D cells

The exposure to progestins produces multiple effects in breast cancer cells which are mediated by the activation of the Progesterone Receptor (PR). The hormone-activated PR translocates to the nucleus where it actively regulates gene transcription. To explore the response to progestins of T47D cells grown in 3D, we measured the level of expression of different transcription factors known to be involved in PR activation in cells cultured in 2D conditions. The levels of PR, ERα and FOXA1 remained similar between 2D and 3D grown cells (Figure Appendix S9A). A slight increase in activated PR phosphorylated at S294 (pPRS294) levels was detected in 3D compared to 2D in the absence and in the presence of 10 nM R5020 (Figure Appendix S9A-B). However, no differences either in the extent of pPRS294 signal induced by hormone or in the percentage of cells responding to the hormonal stimulus was found (Figure Appendix S9B). The pPRS294 signal increased with hormone and its expression was detected homogeneously distributed throughout the cells of the spheroid (Figure S9C, upper panels), similar to the pattern observed for total PR (Figure Appendix S9D).

To globally evaluate the hormone-regulated genes in the spheroids and compare them to the 2D model, we performed RNA-seq experiments. Briefly, T47D cells grown in 2D or 3D were exposed to solvent or to 10 nM R5020 for 6 hours, followed by RNA extraction, mRNA library preparation and massive sequencing. Almost twice as many genes were regulated by hormone in 3D grown cells compared to 2D cells (7,654 *vs* 4,681, log2FC>1, adj p-value <0.01). Up-regulated genes increased around 35%, while down-regulated genes increased by 100 % (Figure 6A). Most of the genes regulated in 3D were not regulated in 2D (4,749 vs 2,961 genes, respectively) (Figure 6B).

**Figure 6.**
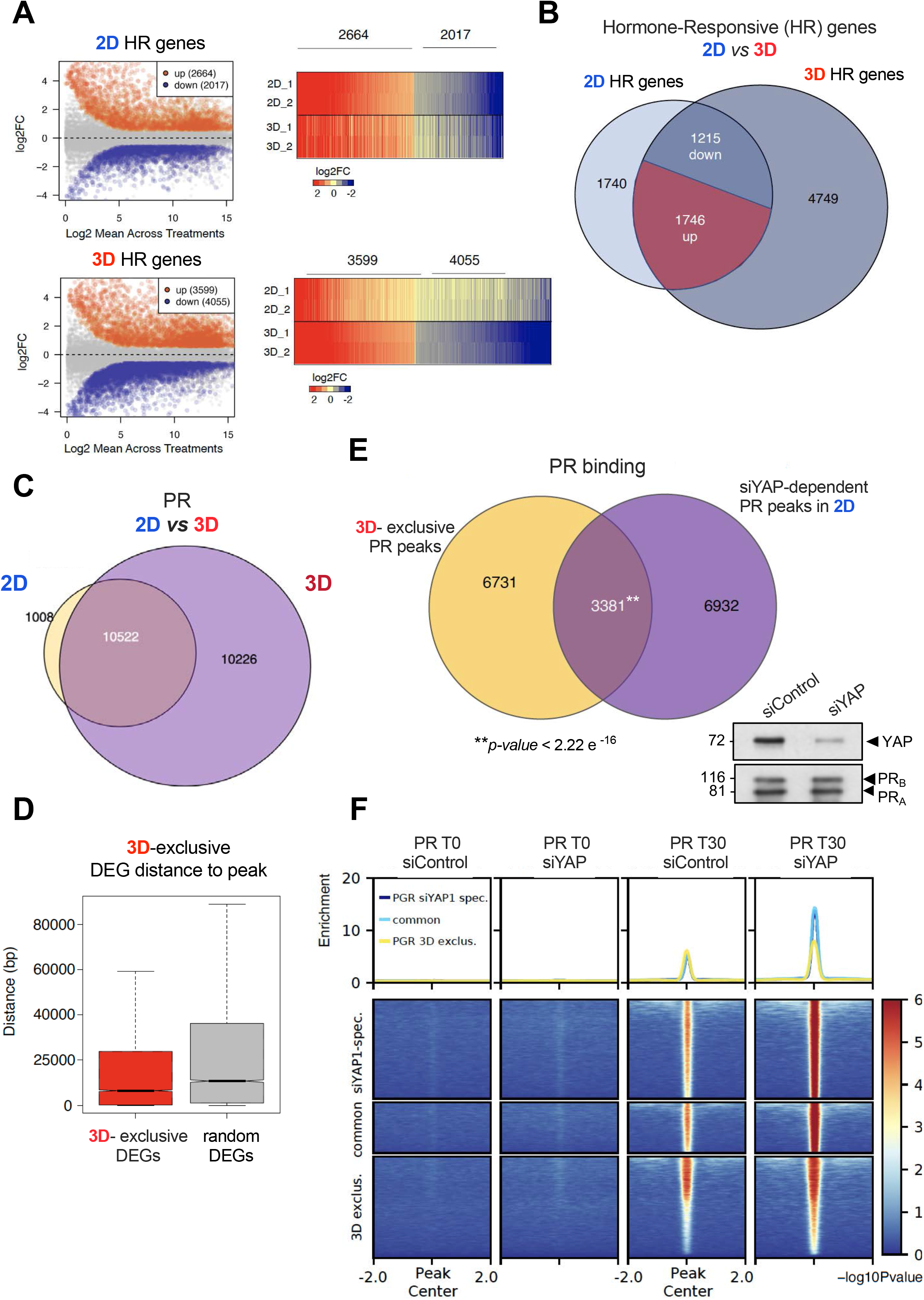
3D-grown cells showed an increased response to the hormone. **A.** T47D cells grown in 2D and 3D conditions in the presence or absence of 10 nM R5020 during 6 h were subjected to RNA-seq assays (log2FC>1, p adj <0.01). The number of genes regulated by hormone in each condition is depicted (left panel). The volcano plot of the distribution of genes regulated in both conditions is shown (right panel). **B.** Venn diagrams of up- and down-regulated genes detected in 2D and 3D cells are shown. **C.** T47D cells grown in 2D and 3D conditions in the presence or absence of 10 nM R5020 during 30 min were subjected to PR ChIP-seq. **D.** Distance between enhancer-associated PRbs (those which overlapped with H3K27ac ChIP-seq signal) to the most proximal differentially expressed gene (3D exclusive). The distance to random genes is used as control. **E.** Venn diagram depicting the overlap of PR binding sites from ChIP-seq experiments performed in siYAP cells treated with hormone and the subset of 3D exclusive PR peaks. Thus, a total of 3,381 PR binding sites of 3D exclusive PR peaks may be a result of YAP displacement detected in the 3D nucleus (p value < 2.2e-16). **F.** Heatmat plots of the PR ChIP-seq experiments performed in siControl and siYAP T47D cells.

Interestingly, among the GO categories regulated by hormone exclusively in 3D we found terms such as *cytoskeleton, actin filament organization, cell growth* and *cell-cell junction* (Appendix Figure S10A-B). These terms were also overrepresented when the extracellular matrix was the only variable incorporated into the analysis (Figure EV1). Thus, when exposed to hormone, T47D breast cancer cells grown as spheroids responded differently to those grown as monolayers.

### PR binding in 2D and 3D grown T47D cells

To explore whether the environmental-dependent changes detected in gene expression are associated with changes on PR binding to the genome, we carried out ChIP-seq experiments of PR in 2D and 3D grown cells exposed to 10 nM R5020.

In line with the increased number of genes regulated in 3D grown cells, we also found more PR binding sites (PRbs) in cells grown as spheroids. Almost all PRbs found in 2D were also present in 3D, but 10,226 new PRbs were exclusively found in 3D grown cells (Figure 6C). The genomic distribution of total PRbs in 3D turned out to be very similar to 2D (Figure EV6A). Compared to 2D exclusive, the distribution slightly changed in the 3D-exclusive PRbs, with less PRbs found in introns and an increased number in promoter, UTR and exons (Figure EV6A).

The majority of the PRbs in 2D grown cells are localized far from the target genes, in enhancers (Ballare et al., 2013). To map the hormone-dependent active enhancers we overlapped 3D-exclusive PRbs (10,226 regions) with H3K27ac peaks obtained from a ChIP-seq performed in 3D grown cells. We measured the distances between these peaks to the nearest significantly regulated gene in 3D grown cells or to random genes. The 3D exclusive genes were significantly closer to 3D PRbs enriched in H3K27ac (enhancers), compared to random genes (Figure 6D). Thus, in 3D grown cells a program is implemented aiming at the regulation of a distinct group of genes that involves the specific binding of PR to 3D-exclusive active enhancers.

### Activation of the Hippo pathway in 3D grown cells impacts in hormone-dependent PR binding

As YAP is a transcriptional coactivator that lack DNA-binding activity, it could act on gene expression *via* interaction with other DNA-binding factors. Reported YAP binding partners include TEAD, p73, Runx2, and the ErbB4 cytoplasmic domain (Li et al., 2010). However, only TEAD has been demonstrated to be important for the growth-promoting function of YAP (Zhao et al., 2008). Interestingly, the 3D exclusive PR binding sites turned out to be enriched in the TEAD DNA binding motif beyond the classical HRE DNA binding motif (Figure EV6B). As in 3D grown cells the Hippo pathway is activated and YAP is phosphorylated in S127 by LATS1 and tagged for degradation in the cytoplasm (Figure 1D), we hypothesize that decreased levels of nuclear YAP would increase the proportion of free TEAD sites, thus, accounting for the enhanced PR binding detected upon hormone in 3D cells.

To address this point, we performed ChIP-seq of PR in siControl and siYAP T47D cells treated with hormone in 2D conditions. We found that 32,7% of PR binding sites that appeared only when YAP is depleted in 2D (10,168 regions) overlap with 3D exclusive PR binding sites (3,381 regions, p value<2.2e^-16^) (Figure 6E). These new sites could regulate a new set of genes associated to an ‘3D spheroid’ condition. Interestingly, PR binding increased globally in the absence of YAP compared to control cells (Figure 6F). According to this model, YAP and PR would compete for binding to 3D exclusive chromatin regions. As YAP does not bind to DNA directly, but rather *via* TEAD, the role of TEAD1, TEAD4 and TAZ in this proposed mechanism should be further elucidated. Recently in our lab, a classification of PR binding sites has been established according to their accessibility at various progestin concentrations. The lowest progestin concentration that allows detection of ligand-dependent PRbs was 50 pM. At this physiological concentration 2,848 PRbs, termed ‘Highly Accessible PR binding sites’, (HAs) were identified (Zaurin et al., 2021) (Figure EV7A).

As HAs constitute essential regulatory elements in hormone-dependent growth of breast cancer cells, we decided to evaluate whether the Hippo pathway also impacted on their function. First, we tested whether HAs-dependent genes required LATS1 activity in 3D grown cells. Our results showed that hormone-dependent regulation of *EGFR*, *STAT5A*, *TIPARP, PGR, BCAS1* and *IGFBP5* was compromised in the presence of TRULI (Figure EV7B). Thus, LATS1 impacts in hormonal response affecting a significant proportion of 3D-exclusive and HAs-associated genes. Unveiling the molecular mechanism by which LATS participates in hormone-dependent gene regulation requires further research.

Therefore, the global impact of the Hippo pathway-through p-LATS activation-in 3D cells could be reflected at least in two ways: i) at basal conditions, via CTCF phosphorylation inducing its displacement and ii) via YAP phosphorylation and inactivation, releasing “hidden” TEAD sites for PR binding allowing regulation of 3D unique genes (Figure EV8). In fact, silencing YAP in 2D partially recapitulate the pattern of PR binding in 3D conditions.

## Discussion

By culturing tubular epithelial breast cancer cells as spheroids (3D) we aimed to explore novel cell signaling pathways, gene networks, transcription factors, chromatin remodelers that might have been overlooked in previous experiments performed in cells cultured as monolayer in plastic dishes (2D). The work reported here on the hormone responsive T47D cell line is a first step in the description and characterization of a more appropriate and physiological in vitro model for breast cancer that should include various epithelial cell types, the supporting fibroblasts and blood cell environment.

Morphologically, T47D cells grown as spheroids in matrigel presented an increase in nuclear volume coupled with a reduction in nuclear diameter (Appendix Figure S1B). These seemingly conflicting alterations might be attributed to the underlying physicochemical modifications that dictate the irregular shapes of the cell nucleus. These processes encompass mechanical forces acting through microtubules, actin filaments, and the osmotic pressure within the cytoplasm (Kim et al., 2015).

It has been widely reported that modifying the stiffness and composition of the used matrix can have an impact on cell growth, cell cycle, differentiation, and the activation of specific signaling pathways, such as the Hippo pathway (Garreta et al., 2019; Uroz et al., 2018). It is known that exposure of cells to a stiffer environment such as plastic implies a force transmission through focal adhesions leading to nuclear flattening and stretching of nuclear pores, reducing their mechanical resistance to molecular transport, increasing YAP nuclear import (Elosegui-Artola et al., 2017), and indirectly affecting gene expression. Conversely, on soft substrates, the nucleus is mechanically uncoupled from the matrix and not submitted to strong forces, inducing a balance between nuclear import and export of YAP through the nuclear pores (Elosegui-Artola et al., 2017) Beyond this mechanical connection between the shape of the nucleus and YAP distribution, we found that 3D cells presented an increased LATS kinase activity which preferentially phosphorylates YAP, reducing its presence in the 3D nucleus (Figure 1D and 5B).

### Impact of 3D growth on chromatin structure and gene regulation

The differential distribution of DAPI nuclear staining in 2D and 3D grown cells pointed to an increased and more structured heterochromatin in 3D grown cells. MNase and ATAC-cleavage experiments provided confirmation that chromatin in 3D cultured cells is characterized by increased compaction, affecting both the larger chromatin fibers and the localized regions where transcription factors interact with nucleosomes. This heightened compaction results in reduced accessibility (Figure 2B-C). In line with these results, we found an increased number of down-regulated genes in 3D grown cells (Figure 1F) and terms related to neurogenesis, lamellipodia and axon guidance are enriched, suggesting higher mobility to migrate in matrigel. In 3D grown cells, we detected protrusions emerging from the cell membrane (pseudopodia) and filopodia extending out from lamellipodia (data not shown), both clear indicators of cell-matrix interactions. These protrusions are associated with cellular sensing mechanisms, involving cell adhesion and cytoskeleton organization strategies (Caswell and Zech, 2018), as supported by the GO terms of the regulated genes.

As T47D cells exhibit an upregulation of genes related to neurogenesis (EV1A), we sought to determine whether this phenomenon was exclusive to breast cancer cells. To address this, we conducted a comparative analysis using non-tumorigenic MCF10A cells cultured as monolayers and in 3D spheres (Maguire et al., 2016). When we examined the genes upregulated in the 3D environment in both cell lines and categorized them by tissue-associated genes, we consistently found an enrichment of brain-associated genes, followed by those associated with the lung, mammary gland, and blood (see Appendix Figure S11A). This suggests that culture conditions have a significant influence on gene expression patterns, rather than being solely dependent on the tumor type itself.

However, upon closer examination of the expression of up-regulated genes in the 3D environment and their clustering, distinctive differences emerged. Notably, in cluster 2, we identified an exclusive overexpression of the term “nervous system development” in T47D cells compared to MCF10A cells (see Appendix Figure S11B). Additionally, we observed significant variations in the expression of genes related to cell and focal adhesion, particularly in cluster 3, which turned out to be distinct between tumor and non-tumoral cells cultured in the 3D environment (see Appendix Figure S11B). Therefore, the distinct expression of neurogenesis-related genes in a 3D environment appears to be associated with a more cancerous behavior. However, further investigations involving a broader range of both tumor and normal cells are required to confirm this observation.

Our results suggest that 3D-gene repression program is driven by a reduction of H3K27ac signal and a concomitant and increase in its trimethylation, while gene activation is led by an increase in H3K18ac, surprisingly, accompanied by H3K9me3 (Figure Appendix S3 and EV2). In this regard, it has been previously reported that H3K9me3 can be found in promoters of repressed genes, as well as at some active genes (Barski et al., 2007).

Additionally, T47D cells grown in 3D exhibit a higher overlap of the nuclear lamina with H3K9me3 and H3K27me3 heterochromatin marks compared to 2D grown cells (data not shown). A tempting hypothesis is that the presence of a more consistent heterochromatin at the Lamina Associated Domains (LADs) might be responsible for the increased gene repression detected in 3D condition. This would imply a 3D-dependent repositioning of genes close to the nuclear lamina, as previously shown in other systems (Reddy et al., 2008). However, more research is required to find the mechanistic and functional basis that support this hypothesis.

### The hippo kinase LATS promotes CTCF displacement from chromatin in 3D grown cells

Despite the changes in chromatin compaction detected by microscopy and nuclease accessibility (Figure 2B-C), when the topology of the genome was assessed through Hi-C experiments, no significant differences in the contact matrices were found at 1 Mb resolution (Figure 3A). However, at higher resolution we detected changes between cells grown in 2D and in 3D at the level of compartments and TAD structure. We found subtle differences indicative of changes in gene repression (more transitions from A to B compartment) (Figure 3B-C). These results along with the identification of the active SEs (Figure EV2D) suggest that the cell identity is maintained in cells grown in 3D (Flyamer et al., 2017; Stadhouders et al., 2019; Vilarrasa-Blasi et al., 2021).

However, we detected moderate changes in genome structure. Compared to cells grown on plastic, cells grown in 3D cells exhibit a higher number of TAD fusions over splits or shifts. Even though the levels of architectural CTCF protein are similar in 2D and 3D grown cells, we detected by ChIP-seq less CTCF bound to chromatin in 3D grown cells (Figure 4A). A possible explanation of this finding could be associated to the increased Hippo pathway activity detected in 3D grown cells, which is translated into a more active p-LATS1 (Figure 5B).

It has been reported that activated p-LATS can phosphorylate CTCF resulting in reduced binding to a subset of genomic binding sites (Luo et al., 2020). In our system CTCF is phosphorylated in 3D grown cells and inhibition or depletion of LATS restores CTCF binding to target regions. Therefore, the Hippo pathway is turned on in 3D grown cells impacting on the cell nucleus through the reduction of CTCF binding, and thus promoting TAD fusions and changes in gene regulation of the neighboring genes. As the dissociation of CTCF we observed in 3D is more global- and not limited to fused TADs-, than previously reported (Luo et al., 2020), the role of p-LATS would be critical and CTCF binding would be more dependent on the Hippo pathway in 3D grown cells. In fact, inhibition of p-LATS by TRULI compromised 3D cell proliferation and reduced the size of the spheroids (Appendix Figure S8A-B).

It is worth to mention, that given the high molecular weight of the CTCF protein (140 KDa) along with the few LATS-dependent phosphorylation sites found (Luo et al., 2020), the effect of TRULI on CTCF phosphorylation could not be detected in Phos-tag gels, even though different conditions were assayed.

It has been reported in breast tissue that LATS activity modulates estrogen receptor (ERα) activity (Lit et al., 2013). More recently, LATS1/2 kinases have been shown to restrict the activity of ERα by binding and promoting its degradation (Britschgi et al., 2017). These studies implicate a nuclear function of LATS kinases in cell lineage commitment and in malignant progression of breast and prostate cancers (Britschgi et al., 2017; Powzaniuk et al., 2004).

Although limited at the level of genome structure, the effect of CTCF depletion detected in 3D had consequences on the activity of 3D-specific genes. Thus, the role of CTCF as a general transcription factor involved in enhancer-promoter looping rather than its function as insulator at TAD borders, would be more relevant in 3D culture cells.

Regarding the uncoupling between displacement of CTCF detected in 3D grown cells and changes in genome architecture, previous work carried out in different systems has shown that CTCF may not be as essential in establishing genome structure/TADs (Barutcu et al., 2018), at least in the mammalian genome, but this is still a matter of debate. In fact, collectively, our findings suggest that the 70% of CTCF displacement detected in 3D chromatin is either not sufficient to change the arrangement of TADs or other factors may be involved and minimize the effect of such substantial depletion. As we previously mentioned, the association of housekeeping genes with boundary regions extends previous studies in yeast and insects (Duan et al., 2010; Ulianov et al., 2016) and suggests that non-CTCF factors may also be involved in insulator/barrier functions in mammalian cells. Although it is a debatable topic, it has been recently reported that very strong loss of CTCF is required (>99%) to detect changes in TADs (Cummings and Rowley, 2022).In fact, additional experiments carried out on compatible systems are needed in order to reach more precise conclusions.

### A more sensitive response to hormone is achieved in 3D cells

Cells grown as spheroids are surrounded by other cells, interact differentially with the ECM and receive nutrients and growth factors in a very heterogeneous manner.

However, in response to hormone, the number of hormone-regulated genes is increased two-fold in 3D grown cells compared to 2D, particularly in repressed genes (2,017 vs 4,055 down-regulated, and 2,664 vs 3,599 up-regulated, in 2D and 3D, respectively). In cells grown in 3D, progestins regulate a group of breast cancer-associated genes that are also regulated in 2D including *CDH10, PGR, CHEK2, LSP1, TERT, SDPR*, but also a new set of 3D-exclusive genes, associated to the ECM including members of the integrin family, Laminins, and *AKT3*.

The response to progestins is more sensitive in 3D grown cells than in cells grown in 2D, as more genes are regulated by hormone. Moreover, the increase in the number of hormone-regulated genes in 3D was accompanied by an increase in the number of PR binding sites detected by ChIP-seq and many of the 3D-exclusive genes are closer to 3D exclusive PR peaks (Figure 6D). Most of these PR binding sites correspond to enhancers, as determined by H3K27 acetylation.

Moreover, LATS1 activity turned out to be essential for high accessible PR binding sites (HAs) (Zaurin et al., 2021) function, as the presence of TRULI compromised hormone-dependent response of HA-associated genes (Figure EV7B). How LATS1 participates in hormonal gene regulation requires further investigation.

Our data support that in 3D grown cells the chromatin is organized differently compared to monolayer cultured 2D cells. DAPI staining, MNase digestion and ATAC-seq assays confirm that the genome is less accessible in 3D (Figure 2), which seems somewhat paradoxical to the increased response to hormone (Figure 6). We found that 3D-activated signals such as the Hippo pathway LATS kinase, impact on the cell nucleus in at least two ways: the LATS kinase phosphorylates both CTCF and YAP promoting the displacement of the first and cytoplasmic retention of the latter. The absence of these two proteins in the 3D nucleus determines the activity of a subset of genes specifically regulated in 3D and in turn, enhances and assures the proper response to hormone (Figure EV8).

In summary, mechanical and chemical signals in 3D grown cells “protect” the nucleus by making it more compact, restricting accessibility to certain regions in the genome and facilitating access to others, thus creating a more sensitive platform for the response to external hormonal cues.

### 3D spheroids: a system that accurately recapitulates the physiological environment of the tumor cell

Cancer cells grown in 3D culture systems exhibit physiologically relevant cell-cell and cell-matrix interactions, gene expression and signaling pathway profiles, heterogeneity and structural complexity that reflect *in vivo* tumors (Nath and Devi, 2016). Actually, the non-cellular components of the tumor microenvironment (TME), those that are preserved in spheroids, as the extracellular matrix (ECM), growth factors, cytokines, and chemokines, play a significant role in cancer progression (Paszek et al., 2005). We found that breast cancer 3D spheroids showed high expression of key genes involved in tumor growth, as well as LATS1-dependent nuclear receptor binding features shared with PDXs but absent in 2D cells (Figures EV1B and EV6). In line with this, LATS1 has been linked to cancer cell plasticity and increased resistance to hormone therapy in breast tumors (Furth and Aylon, 2017).

To assess whether our 3D system recapitulate the resistance of breast cancer cells to CDK4/6 inhibitors and endocrine therapy, which is associated with low FAT1 levels and Hippo pathway activation, we conducted a comparative analysis of gene expression data. Specifically, we compared the gene expression profiles of our 3D breast cancer cell model with those from a previous study (Li et al., 2018). Our analysis revealed that cells resistant to CDK4/6 inhibitors and endocrine therapy exhibited reduced levels of FAT1 and demonstrated activation of the Hippo pathway, which matched the characteristics observed in our 3D-grown cells (as depicted in Appendix Figure S12). However, the resistant cells had lower progesterone receptor (PR) status (Appendix Figure S12A), leading to decreased expression of genes controlled by progestin, both in terms of activation and repression (Appendix Figure S12D). Thus, although our 3D model partially replicated the characteristics of resistant cells, it did not entirely reproduce the diminished hormonal response observed in cells lacking FAT1 expression (Appendix Figure S12E). In fact, in our 3D model, this response was enhanced (Figure 6A).

Regarding the heterogeneity, gene expression analysis showed that 3D spheroids are enriched in terms associated to nervous system development compared to 2D cells (Figure 1F and EV1A). In fact, increasing evidence suggests that the nervous system itself, as well as neurotransmitters and neuropeptides present in the tumor microenvironment, play a role in orchestrating tumor progression (Fernandez-Nogueira et al., 2016). Thus, 3D cells accurately recapitulate the intratumor heterogeneity facilitating tumor progression and fostering the adaptation and survival of the different tumor cells to the different microenvironments in which a tumor resides.

Our results highlight the importance of the 3D system as a reliable model lacking cross-species incompatibilities for breast cancer study.

## Experimental procedures

### Cell Culture and hormone treatments

T47D breast cancer cells were routinely grown in RPMI 1640 medium supplemented with 10% FBS, 2 mM L-glutamine, 100 U/ml penicillin and 100μg/ml streptomycin. BT-474, ZR-75 and MCF-10A cells were obtained from ATCC and cultured in 2D and 3D in the recommended medium.

For the experiments, cells were plated in RPMI medium without phenol red supplemented with 10% dextran-coated charcoal treated FBS (DCC/FBS) and 48 h later medium was replaced by fresh medium without serum. After 24h in serum-free conditions, cells were incubated with 10 nM R5020 for different times at 37°C.

MCF-10A cells were routinely grown in DMEM/F12 medium with 5% Horse Serum, 20 ng/ml EGF, 100 U/ml penicillin and 100mg/ml streptomycin, 0.5 mg/mL Hydrocortisone, 100 ng/ml Cholera Toxin and 10 mg/ml Insulin.

### 3D cell culture on Matrigel

Prechilled p60 plates were coated with a thin layer of Matrigel (Corning Life Sciences) and then incubated for 20–30 min at 37°C to allow polymerization without over drying. 200,00 cells trypsinized cells were resuspended and plated on-top of the Matrigel without disrupting the capsule. Culture was maintained for 10 days changing the medium every 2-3 days. For hormonal induction on the 3D spheroids, T47D cells were plated on top of phenol red-free Matrigel and after 7 days of culture medium was replaced by RPMI medium without phenol red supplemented with 10% dextran-coated charcoal treated FBS (DCC/FBS) and 48 h later medium was replaced by fresh medium without serum. After 24h in serum-free conditions, cells were incubated with 10 nM R5020 for different times at 37°C.

*Spheroids extraction from 3D matrix:* Dishes containing the spheroids grown for 10 days were rinsed twice with PBS followed by addition of 3 ml ice-cold PBS-EDTA. Matrigel including the 3D spheroids was carefully detached with a plastic scraper and left on ice for 30 min to allow complete depolymerization of the gel. The liquid solution was transferred to a conical tube, centrifuged for 5 min at 112 x *g* and rinsed twice with 0.5 volume of PBS-EDTA. The cell pellet was then ready for further processing.

### Chromatin Immunoprecipitation (ChIP) in 2D and 3D cultured cells

ChIP assays were performed as described (Strutt and Paro, 1999) using anti-PR (H190 SC-7208, Santa Cruz,); anti-RNApol II (#2629, Cell Signaling); anti-CTCF (07-729, Merck); anti-H3K27ac (ab4729, Abcam); anti-H3K18ac (#39693, Active Motif). Quantification of chromatin immunoprecipitation was performed by real time PCR using Roche Lightcycler (Roche). The fold enrichment of target sequence in the immunoprecipitated (IP) compared to input (Ref) fractions was calculated using the comparative Ct (the number of cycles required to reach a threshold concentration) method with the equation 2 ^Ct(IP)-Ct(Ref)^. Each of these values were corrected by the human β-globin gene and referred as relative abundance over time zero. Primers sequences for target regions are available on request.

### RNA interference experiments

Stable LATS depleted T47D cells were generated by using lentiviral shRNAs obtained from Sigma (MISSION) shRNA Lentiviral Transduction Particles: TRCN0000001777_LATS and TRCN0000001779_LATS.

### Cell Proliferation

T47D cells (1×10^3^) were plated in non-transparent-walled 96-well plate in RPMI medium or in charcolized-medium in the presence or absence of 10nM R5020. The TiterGlo reagent (Promega) was added, cells were then incubated for 2 min at RT with agitation, followed by 10 min incubation at RT. Bioluminescence was detected in a Barthold luminometer system allowing 0.25 second per well. The experiments were performed in quintuplicate.

### Flow Cytometry

T47D cells were plated into duplicate wells of six-well plastic dishes and preincubated as described. After 24h 10nM R5020 was added. Cells were harvested at the start of treatment (control, zero time) and after 24h of hormone addition. The cell suspension was pelleted, stained with propidium iodide and treated with ribonuclease (RNase). Samples were cooled to 4°C, and 10,000 cells were analyzed on a BD FACSCanto analyser flow cytometer.

### Live/dead assay

To visualize the number of viable cells in a 3D spheroid the LIVE/DEAD™ Cell Imaging Kit was used according to the manufacturer protocol.

### Immunofluorescence

For immunostaining, 1×10^3^ cells per well were seeded on Matrigel precoated 8-well LabTek (10 mm). 2D and 3D cells grown for 3 and 10 days respectively, were washed two times with PBS and fixed for 10 min with 4 % fresh Paraformaldehyde (PFA), before being permeabilized during 30 min with 0.5% Triton X-100 in PBS. Samples were then blocked during 1.5 h with 5 % BSA solution. Then, the cells were incubated o/n at 4°C with the corresponding antibody diluted in the IF solution.

The following day, cells were washed three consecutive times with IF solution during 10 min each. The samples were then incubated during 1h at 4 °C with the secondary antibody in a humid dark chamber.

The secondary antibodies used were: AlexaFluor 488 anti-rabbit IgG (1:1000; raised in donkey) and AlexaFluor 546 anti-mouse (1:1000; raised in goat. After the incubation with the secondary antibody, cells were washed once with IF solution, incubated during 10 min with 0.1 mg/ml DAPI in PBS and then washed with PBS three times, before mounting in Mowiol 4-88 Mounting Medium for imaging. Images were collected sequentially on a Leica SP8-STED confocal laser-scanning microscope using the software Leica Application Suite X. All collected images conserved an optical thickness of 0.25 µm. Image analysis for marker distribution, quantification and colocalization were performed using ImageJ (Schindelin et al., 2012).

### Preparation of cell extracts and western blot

Matrigel-free cell pellets were collected and washed two times with PBS-EDTA and lysed with RIPA buffer followed by incubation at 95°C for 10 min. For western blotting, pellets were centrifuged and quantified by Bradford before being loaded and run in SDS-acrylamide gels. The following antibodies were used: CTCF (07-729, Millipore); ERα (H20, Santa Cruz); PR (H190, Santa Cruz); FAK (PTK2) (3285, Cell Signaling); LATS (3477, Cell Signaling); LATS-S909 (9157, Cell Signaling); pFAKT397 (ab81298, Abcam); pPRs294 (ab61785, Abcam); pYAP127 (4911, Cell Signaling) and YAP (sc-101199, Santa Cruz).

### Phos-tag gels

Whole cell lysates were separated on a 5% SDS/PAGE containing 20 μm Phos-tag (NARD Institute), followed by western blotting with anti-CTCF antibody.

### MNase digestion

2D and 3D grown T47D cells were washed once with PBS, collected in 2 ml cold PBS + PIC and centrifuged 5 min at 900 x *g* at 4°C. The cell pellet was then gently resuspended in 50 μl RBS buffer (10 mM Tris-HCl pH 7.4, 10 mM NaCl, 3 mM CaCl2.) followed by addition of 1.3 ml RBS buffer + 0.1 % NP40. Cells were centrifuged again for 10 min at 500 x *g* at 4°C and the nuclei were then resuspended in RBS buffer for counting. An amount of 600,000 nuclei obtained from 2D and 3D grown cells were treated with 0, 30, 45, 90 Units of MNase to obtain differential digestion patterns. The MNase reaction was carried out in 500 μl final volume reaction. The nuclei were then incubated for 2 min at 37°C. The reaction was stopped with 40 mM EDTA 0.5 M and then treated with RNAseA 10 mg/ ml for 30 min at 37°C and Proteinase K (1.2 μg/ μl) for 1 h at 45°C. The DNA was purified with Phenol/Chloroform and 600 ng of material was loaded in a 1.2 % agarose gel.

### RNA-seq

RNA was extracted from T47D cells grown in 2D and 3D conditions in RPMI 1640 medium. To evaluate the effect of the hormone, 2D and 3D T47D cells were treated or not for 6hs with 10nM R5020 and submitted to massive sequencing using the Solexa Genome Analyzer. The protocol followed to analyze the RNA-seq data can be found in the Supplementary methods section.

### ChIP-seq

ChIP-DNA was purified and subjected to deep sequencing using the Solexa Genome Analyzer (Illumina, San Diego, CA). The protocol followed to analyze the ChIP-seq data can be found in Supplementary methods section.

### ATAC-seq

ATAC experiments were performed as described (Buenrostro et al., 2013) using nuclei obtained from 2D or 3D grown T47D cells. Extended bioinformatics methods can be found in the Supplementary information.

### Hi-C experiments

Hi-C libraries were generated from 2D and 3D grown T47D cells treated or not with R5020 for 60 min according to the previously published Hi-C protocol with minor adaptations (Lieberman-Aiden et al., 2009). Hi-C libraries were generated independently in both conditions using HindIII and NcoI restriction enzymes. Hi-C libraries were controlled for quality and sequenced on an Illumina Hiseq2000 sequencer. The Illumina Hi-seq paired-end reads were processed by aligning to the reference human genome (GRCh37/hg19) using BWA.

## Acknowledgements

The experimental work was supported by grants from the Departament d’Innovació Universitat i Empresa (DIUiE), the Spanish Ministry of Economy and Competitiveness (SAF2016-75006-P and PID2019-105173RB-I00) and Consejo Superior de Investigaciones Científicas (Ref# 201820I131), ‘Centro de Excelencia Severo Ochoa 2013-2017’, SEV-2012-2018 and ERC Synergy Grant “4DGenome” nr: 609989. MAM-R acknowledges support by the Spanish Ministerio de Ciencia e Innovación (PID2020-115696RB-I00).

## EXTENDED BIOINFORMATICS METHODS

ChIP-seq and ATAC-seq Peak Calling: Analysis of sequence data was carried out as previously described (2) with minor modifications. Reads were aligned to the hg38 human genome reference sequence (GRCh38) using Bowtie (3) and aligning parameters of uniqueness (-S –m1 –v2 –t –q). p values for the significance of ChIP-seq counts compared to input DNA were calculated as described (4) using a threshold of 10-8 and a false discovery rate (FDR) < 1%.

### ChIP-seq Downstream Analysis

Average ChIP-seq signals of 50 bp windows around 1 kb upstream and downstream of annotated TSSs were calculated using the cis-regulatory annotation system (CEAS) (5).

### ATAC-seq Downstream Analysis

PoissP wiggle generate from peaks calling (above) were used with Sitepro (5) to calculate the average nucleosome signal across genome coordinates of selected BED.

### Motif analysis

Homer motif analysis (6) was conducted on all PRBs as represented in figure 2E). FIMO and SpaMo (3,5) were used respectively to calculate the frequency and the precise location of PR, FOXA1 and ERa corresponding binding motifs (HRE, FOXA1 and ERE). Matrixes used for each factor (PGR/MA0727.1) (ERS1/MA0112.2), (FOXA1/MA0148.2) are from Jaspar database (1).

### RNA-seq

#### RNA extraction, RNA-seq library preparation, and qRT-PCR

RNA was isolated from cells cultured in 2D monolayer or as 3D spheroids with TRIzol reagent (Ambion), ethanol precipitated, and dissolved in sterile water. RNA concentration was measured with a Qubit fluorometer and RNA subjected to Bioanalyzer for quality control. Libraries were prepared using 1 μg of polyA+ RNA by PCR amplification of cDNA with bar-coded primers using the Illumina TruSeq kit at the CRG Genomic Facility. Libraries were sequenced using Illumina HIseq-2500 to obtain pair-ended (PE) 100-base-long reads.

For gene expression analysis, RNA (250 ng) was subjected to cDNA synthesis using the qScript cDNA Synthesis kit (Quanta Biosciences). qPCR was carried out using the LightCycler FastStart DNA Master SYBR Green I kit (Roche), and specific primers selected from the list of available designed primers at Primer Bank (https://pga.mgh.harvard.edu/primerbank) (7). As reference gene, GAPDH was used.

#### RNA-seq Pipeline and Differential Gene Expression Analysis

Sequencing adapters and low-quality ends were trimmed from the reads using Trimmomatic, using the parameters values recommended (8) and elsewhere (https://goo.gl/VzoqQq) (trimmomatic PE raw_fastq trimmed_fastq ILLUMINACLIP:TruSeq3-PE.fa:2:30:12:1:true LEADING:3 TRAILING:3 MAsXINFO:50:0.999 MINLEN:36). The trimmed reads were aligned to GRCh38 (9) using STAR (10).

First, the genome index files for STAR were generated with: star–runMode genomeGenerate–genomeDir GENOME_DIR–genomeFastaFiles genome_fasta– runThreadN slots–sjdbOverhang read_length–sjdbGTFfile sjdb–outFileNamePrefix GENOME_DIR/Where genome_fasta is the FASTA file containing the GRCh38 sequence downloaded from the University of California Santa Cruz (UCSC) Genome Browser, excluding the random scaffolds and the alternative haplotypes; and sjdb is the GTF file with the GENCODE’s V24 annotation.

Second, trimmed reads were aligned to the indexed genome with: star–genomeDir GENOME_DIR/–genomeLoad NoSharedMemory–runThreadN slots–outFilterType “BySJout”–outFilterMultimapNmax 20–alignSJoverhangMin 8–alignSJDBoverhangMin 1–outFilterMismatchNmax 999–outFilterMismatchNoverLmax 0.04–alignIntronMin 20– alignIntronMax 1000000–alignMatesGapMax 1000000–readFilesIn read1 read2– outSAMtype BAM SortedByCoordinate–outTmpDir TMP_DIR/–outFileNamePrefix ODIR1/$sample_id.–outWigType bedGraph–readFilesCommand zcat.

Differences in gene expression were calculated by using a DESeq.R script for RNA analysis (available for download at Mendeley http://dx.doi.org/10.17632/mzjf96t3gc.5). Genes with fold change (FC).} 1.5 (p value < 0.05; FDR < 0.01) were considered as significantly regulated. Sitepro profiles were generated with the script, provided in the CEAS package.

### *In situ* Hi-C library preparation

*In situ* Hi-C was performed as previously described (11) with the following modifications:

i. two million cells obtained from T47D cells cultured in 2D monolayer or as 3D spheroids were used as starting material; (ii) chromatin was initially digested with 100 U MboI (New England BioLabs) for 2 h, and then another 100 U (2 h incubation) and a final 100 U were added before overnight incubation; (iii) before fill-in with bio-dATP, nuclei were pelleted and resuspended in fresh 1 NEB2 buffer; (iv) ligation was performed overnight at 24°C with 10,000 cohesive end units per reaction; (v) de-cross-linked and purified DNA was sonicated to an average size of 300–400 bp with a Bioruptor Pico (Diagenode; seven cycles of 20 sec on and 60 sec off); (vi) DNA fragment-size selection was performed only after final library amplification; (vii) library preparation was performed with an NEBNext DNA Library Prep Kit (New England BioLabs) with 3 μl NEBNext adaptor in the ligation step; (viii) libraries were amplified for 8–12 cycles with Herculase II Fusion DNA Polymerase (Agilent) and were purified/size-selected with Agencourt AMPure XP beads (>200 bp). Hi-C library quality was assessed through ClaI digestion and low-coverage sequencing on an Illumina NextSeq500 instrument, after which every technical replicate (n=2) of each biological replicate (n=2) was sequenced at high coverage on an Illumina HiSeq2500 instrument. Data from technical replicates were pooled for downstream analysis. We sequenced >18 billion reads in total to obtain 0.78–1.21 billion valid interactions per time point per biological replicate.

### *In situ* Hi-C data processing and normalization

We processed Hi-C data by using an in-house pipeline based on TADbit (12). First, the quality of the reads was checked with FastQC to discard problematic samples and detect systematic artifacts. Trimmomatic (8) with the recommended parameters for paired-end reads was used to remove adaptor sequences and poor-quality reads (ILLUMINACLIP:TruSeq3-PE.fa:2:30:12:1:true; LEADING:3; TRAILING:3; MAsXINFO:targetLength:0.999; and MINLEN:36).

For mapping, a fragment-based strategy implemented in TADbit was used, which was similar to previously published protocols (13). Briefly, each side of the sequenced read was mapped in full length to the reference genome (GRCh38). After this step, if a read was not uniquely mapped, we assumed that the read was chimeric, owing to ligation of several DNA fragments. We next searched for ligation sites, discarding those reads in which no ligation site was found. The remaining reads were split as often as ligation sites were found.

Individual split read fragments were then mapped independently. These steps were repeated for each read in the input FASTQ files. Multiple fragments from a single uniquely mapped read resulted in a number of contacts identical to the number of possible pairs between the fragments. For example, if a single read was mapped through three fragments, a total of three contacts (all-versus-all) was represented in the final contact matrix. We used the TADbit filtering module to remove non-informative contacts and to create contact matrices. The different categories of filtered reads applied were:

1. Self-circle: reads coming from a single restriction enzyme (RE) fragment and pointing to the outside.
2. Dangling end: reads coming from a single RE fragment and pointing to the inside.
3. Error: reads coming from a single RE fragment and pointing in the same direction
4. Extra dangling end: reads coming from different RE fragments but that were sufficiently close and point to the inside; the distance threshold used was left to 500 bp (default), which was between percentiles 95 and 99 of average fragment lengths.
5. Duplicated: the combination of the start positions and directions of the reads was repeated, thus suggesting a PCR artifact; this filter removed only extra copies of the original pair.
6. Random breaks: the start position of one of the reads was too far from RE cutting site, possibly because of non-canonical enzymatic activity or random physical breaks; the threshold was set to 750 bp (default), >percentile 99.9.

From the resulting contact matrices, low-quality bins (those presenting low contact numbers) were removed, as implemented in TADbit’s ‘filter columns’ routine. A single round of ICE normalization (14), also known as ‘vanilla’ normalization (16), was performed. That is, each cell in the Hi-C matrix was divided by the product of the interactions in its columns and the interactions in its row. Finally, all matrices were corrected to achieve an average content of one interaction per cell.

### Identification of subnuclear compartments and topologically associated domains (TADs)

To segment the genome into A/B compartments, normalized Hi-C matrices at 100-kb resolution were corrected for decay as previously described, by grouping diagonals when the signal-to-noise ratio was below 0.05 (11). Corrected matrices were then split into chromosomal matrices and transformed into correlation matrices by using the Pearson product-moment correlation.

Normalized contacts matrices at 20-kb resolution were used to define TADs, and for visualization purposes, through a previously described method with default parameters (15,16). First, for each bin, an insulation index was obtained on the basis of the number of contacts between bins on each side of a given bin. Differences in the insulation index between both sides of the bin were computed, and borders were called, searching for minima within the insulation index. The insulation score of each border was determined as previously described (16), by using the difference in the delta vector between the local maximum to the left and the local minimum to the right of the boundary bin. This procedure resulted in a set of borders for each time point and replicate. To obtain a set of consensus borders along the time course, we proceeded in two steps: (i) merging borders of replicates and overlapping merged borders (that is, for each pair of replicates, we expanded the borders one bin on each side and kept only those borders present in both replicates as merged borders) and (ii) further expanding two extra bins (100 kb) on each side and determining the overlap to obtain a consensus set of borders common to any pair of time points.

**EV1**: Mutant or Translocated Estrogen Receptor Alpha (MOTERA) signature heatmap for our RNA-seq experiments of 3D versus 2D gene expression was generated recovering the log2 ratio (3Dvs2D) for each of the 24 genes belonging to the signature. Cluster 3.0 was used to generate the CDT file loaded on Java Tree View for heatmap visualization. Genes were ranked from the highest expressed to the lowest expressed.

**EV3**: Super-enhancers (SE) were calculated using Richard Young ROSE algorithm (https://bitbucket.org/young_computation/rose/src/master/) by applying H3K27ac data from ChIP-seq experiments in 2D and 3D conditions. Default parameter were used for SE computation. Gene association was done by using bedtools closest-features with the closest genes being the more highly expressed. A and B compartments were called using Homer (6) Sub-nuclear Compartment Analysis (PCA/Clustering).

**EV4**: Homer motif analysis (6) was conducted on 2D-and 3-D-specific ATAC-seq peaks using default parameters. Word clouds profile of transcription factor binding was generated using Toolkit analysis (http://dbtoolkit.cistrome.org/). The table reporting the GIGGLE score for each of the factor to be bound at these ATAC 2D and 3D peaks (retrieved from Toolkit) was used to extrapolate the frequency of the factor bound and plotted using text mining/word cloud using R package.

**Appendix Figure S5**: **A.** RAD21 ChIP-seq were analyzed as described in Material and Methods. For each ChIP-seq deepTools (Ramirez et al., 2014) was used to generate the meta-TAD analysis reporting the log10 of all RAD21 ChIP-seq normalized tags around and within the TAD fused. A random set of TADs was used as a control. Boxplots have been generate using data from panel A using R boxplot package. Two-way ANOVA test was used to calculate statistical significance.

**Appendix Figure S7**: **A.** *Drosophila M.* spike in CTCF ChIP-seq. Total tags form Drosophila M. retrieved in the ChIP experiment were counted and reported as a bar graph, for two biological replicates. Genome browser view of chromosome 2L form dm6 build was generated after H2av tags were compared with the input control for dm6. **B.** The IGV genome browser snapshot clear shows peaks of enrichment attesting the good immunoprecipitation of the H2av. For each CTCF sample we calculated the 3D vs 2D tag normalization factor given the spike-in tags retrieve in the H2av immunoprecipitation (as reported in the table). This factor was used to normalized the corresponding CTCF ChIP experiments (by applying this factor to the tag counts of the bam file). The resulted spike-in-normalized bam were used to downstream peak calling as described in material and methods. **C.** Heatmaps of Spike-in normalized CTCF ChIP-seq were generated using deepTools. From two biological replicates we generated one unique file by merging the two file and considering only the common peaks. We then used the 2D CTCF peaks to monitor the decrease of CTCF upon 3D growth in the presence or absence of TRULI (Panel C). We plotted heatmaps of enrichment using deepTools (Ramirez et al., 2014).

**Appendix Figure S11: A.** Comparison T47D vs MCF10A (Maguire et al., 2016). Gene up-regulated in T47D 3D and genes up-regulated in spheroid versus plastic MCF10A were analyzed for tissue-specific enrichment by employing SEdb2.0 (https://bio.liclab.net/sedb/analysis_gene.php). **B.** Heatmap of T47D 3D growth-regulated genes, compared to MCF10A spheroid versus plastic. Only 50% of the total T47D was detectable in MFC10A with various degree of expression inversions as in cluster (K) 2 and K3. Clusters K1 and K4 show parallel gene expression for both up- and down-regulation between the two cell types. Table of Gene Ontology (GO) for the cluster reported in the heatmap.

**Appendix Figure S12**: **A.** Expression levels of FAT1 and PGR in parental and FAT1 KO (CR) cells (Li et al., 2018). **B.** Expression of FAT1 in our T47D cells both growth in 2D and 3D. For both conditions the four replicate experiments levels are reported. **C.** Genome browser view of FAT1 and PGR gene expression levels. **D.** Boxplot comparison of hormone-dependent up- and down-regulated genes in in both 2D and 3D with and without R5020 treatment. The expression levels of these groups are compared also with the same genes but in the RNA-seq from FAT1 KO cells (Li et al., 2018). **E.** Heatmap of expression of important hormone regulated genes.

## Expanded view

**EV1.**
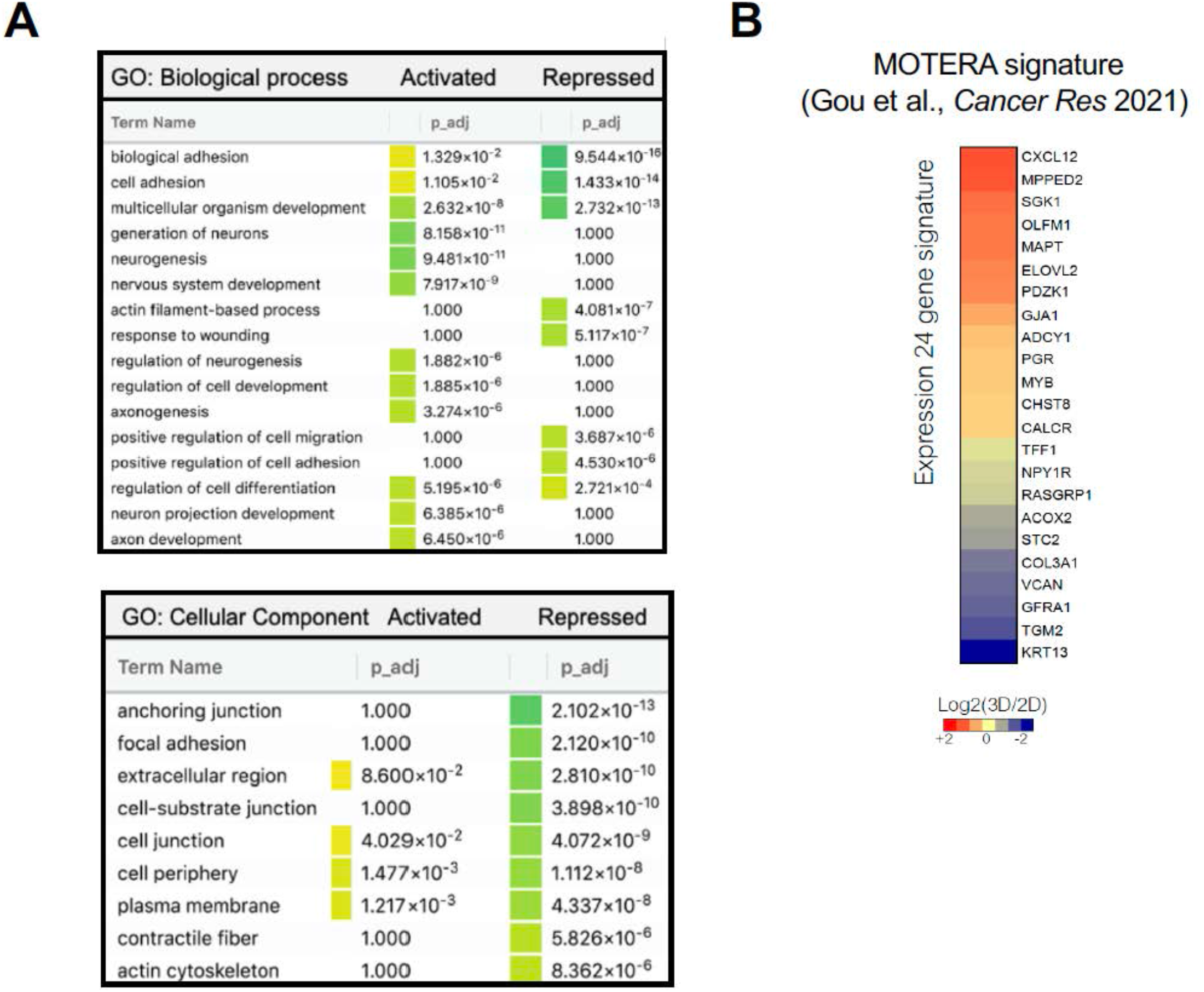
Gene Ontology of Biological Process (BP) and Cellular Component (CC) differentially regulated in 3D cells. **A.** Using DEA results, GO was performed for Biological Process (upper panel) and Cellular component (lower panel). The main terms enriched are associated with cell adhesion processes, cell structure and regulation of neurogenesis. **B.** Mutant or Translocated Estrogen Receptor Alpha (MOTERA) signature heatmap for our RNA-seq experiments of 3D versus 2D gene expression was generated recovering the log2 ratio (3Dvs2D) for each of the 24 genes belonging to the signature. Cluster 3.0 was used to generate the CDT file loaded on Java Tree View for heatmap visualization. Genes were ranked from the highest expressed to the lowest expressed.

**EV2.**
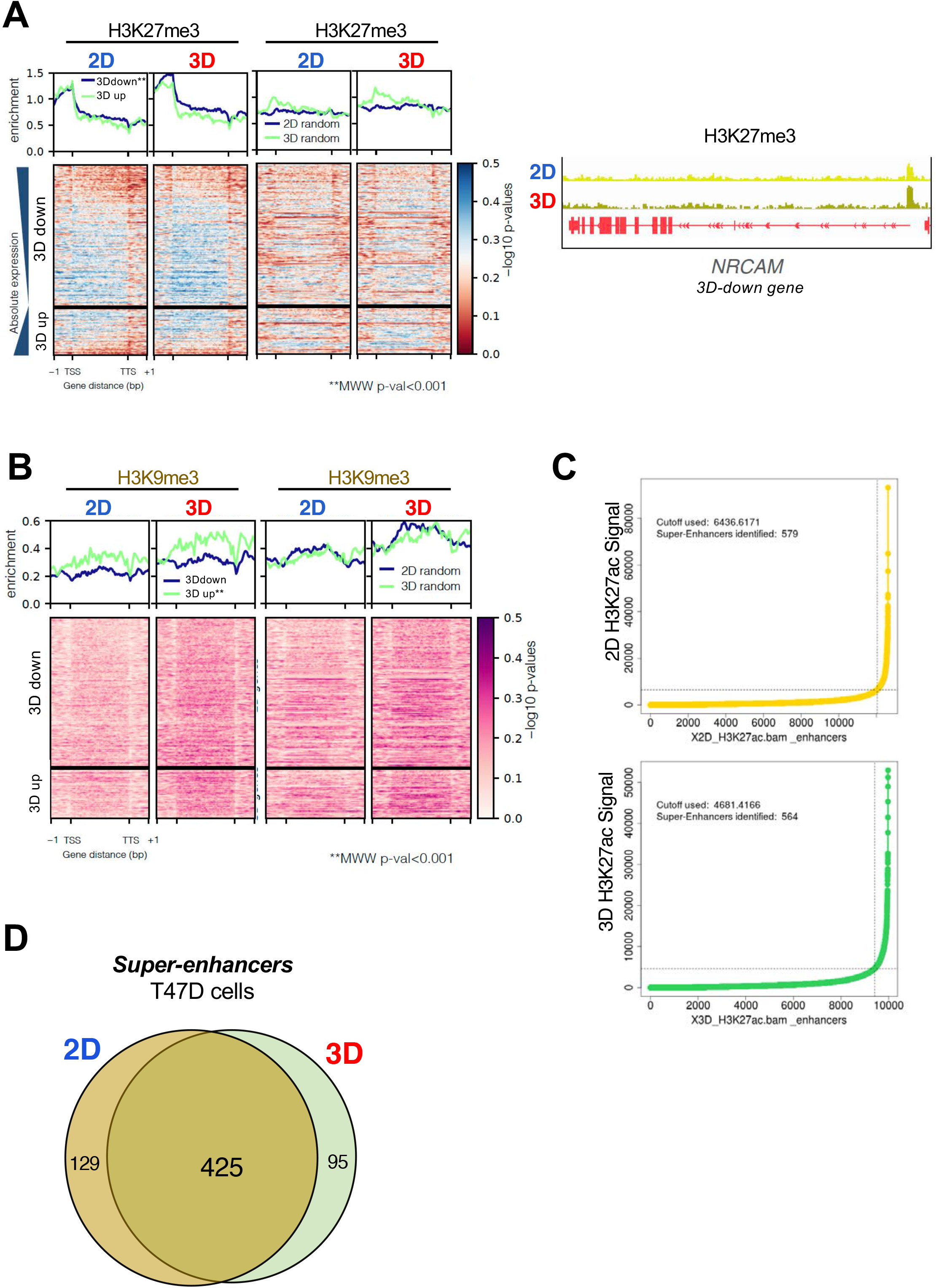
Profiles of H3K27me3, H3K9me3 and super-enhancers detected in 2D and 3D T47D breast cancer cells. **A.** The H3K27me3 profiles in 3D repressed and activated genes (blue and green lines, respectively) obtained in both conditions (first and second panels, from the left) is shown. Third and fourth panel from the left: profiles of H3K27me3 in 2D and 3D random genes. *Right panel:* Genome browser view of H3K27me3 ChIP-seq data in the *NRCAM* gene. **B.** The H3K9me3 profiles in 3D repressed and activated genes (blue and green lines, respectively) obtained in both conditions (first and second panels, from the left) is depicted. Third and fourth panels from the left: profiles of H3K9me3 in 2D and 3D random genes. **C.** The enrichment of the H3K27ac signal in super-enhancers obtained from 2D and 3D cells is shown. **D**. Venn diagram corresponding to the super-enhancers detected in T47D cells grown in 2D and 3D conditions.

**EV3.**
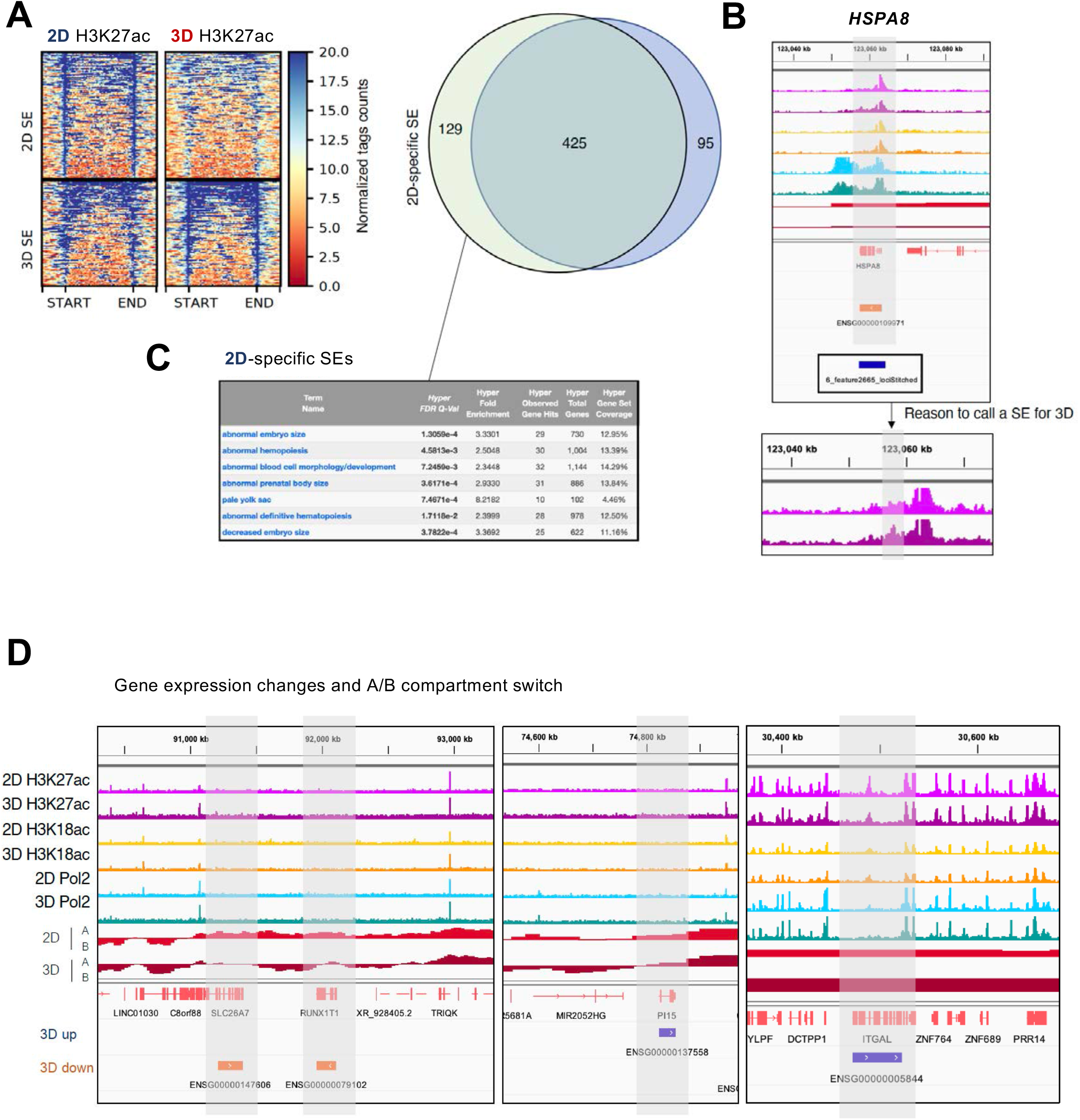
Characterization of super-enhancers identified in 2D and 3D cells. **A.** The enrichment of the H3K27ac signal in super-enhancers obtained from 2D and 3D cells is shown. Venn diagram corresponding to the super-enhancers detected in T47D cells grown in 2D and 3D conditions. By using a proximity-based script (Hnisz et al., 2013), we found 129 and 95 genes associated to 2D and 3D SEs, respectively (right panel). In the case of genes exclusively regulated in the 3D condition by SEs, many of them appear to be artifacts, as illustrated with the HSPA8 gene (**B**). **C.** The 2D-exclusive genes are related to terms like abnormal embryo size, abnormal development, and hematopoiesis. **D.** Snapshots from the genome browser illustrating the transitions for two 3D down-regulated genes, SLC26A7 and RUNX1T1, and two 3D up-regulated genes, PI15 and ITGAL are shown.

**EV4.**
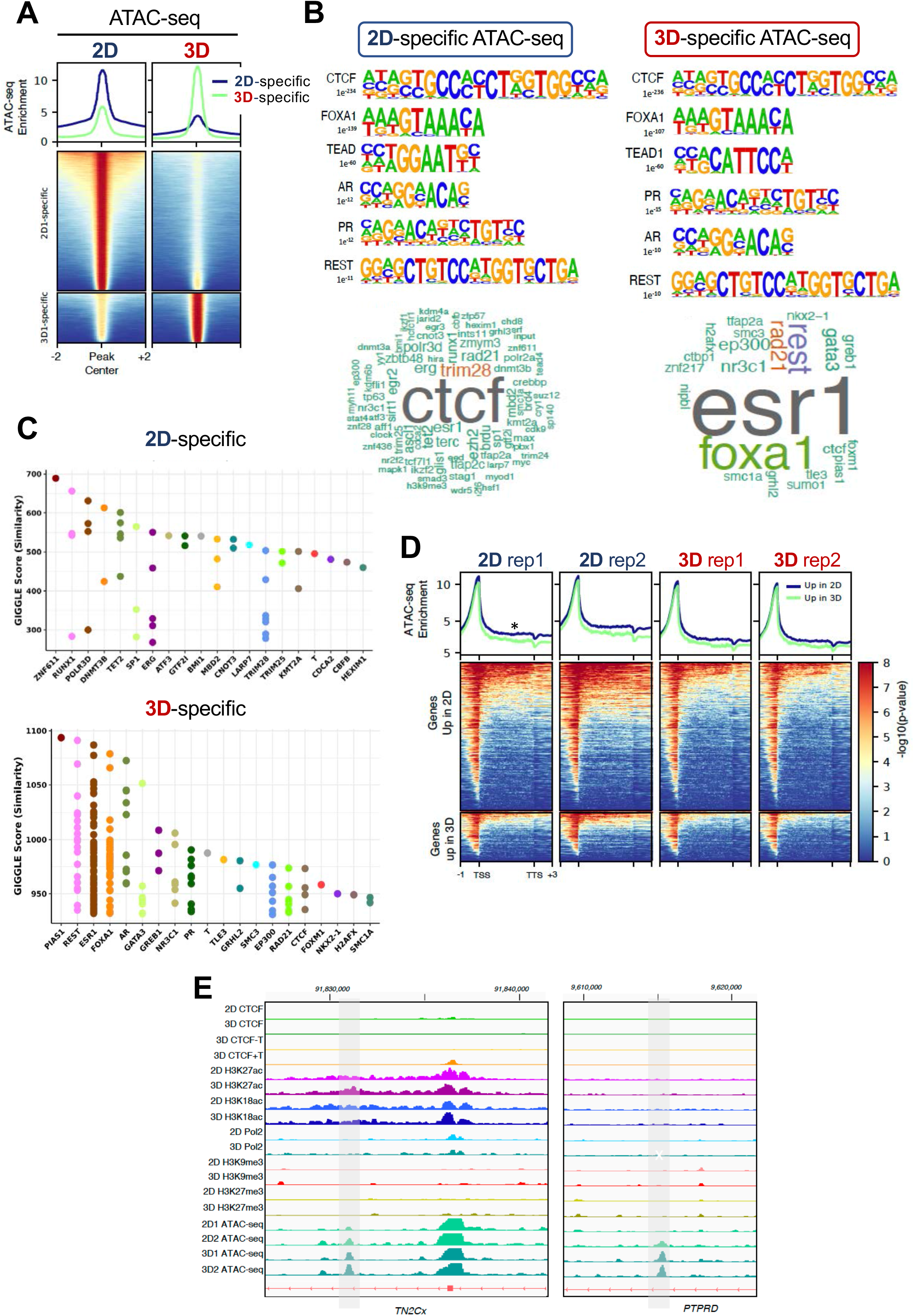
Regions more accessible in 3D are enriched in estrogenic signalling. **A.** Heatmaps of ATAC-seq data performed in 2D and 3D T47D cells. The exclusive regions belonging to each condition is highlighted. **B.** Homer motif analysis of 2D and 3D-exclusive ATAC-seq regions (upper panel). When the same regions are contrasted with available ChIP-seq data, the CTCF and ESR1 terms appear enriched (bottom panels). **C.** Giggle score of the data presented in panel B. **D.** Heatmaps of the ATAC-seq signal around 2D and 3D up genes. **E.** Snapshot of the genome browser around TN2Cx and PTPRD genes showing the profiles of H3K27ac, RNAPol2, H3K9me3, H3K27me3 and ATAC-seq.

**EV5.**
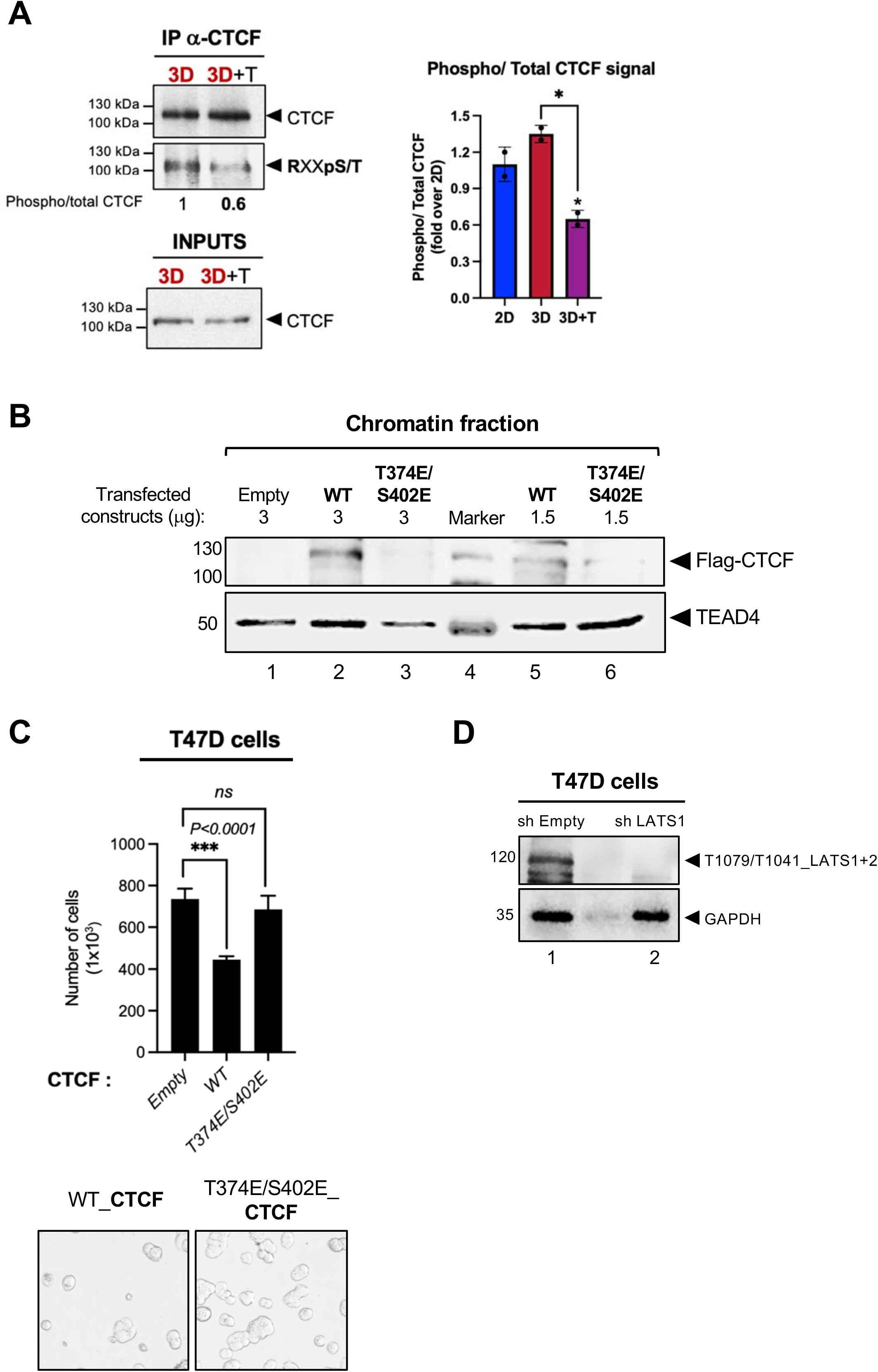
CTCF phosphorylation is reduced in the presence of LATS inhibitor. **A.** 2D, 3D and 3D treated with TRULI T47D cells were cultured and subjected to immunoprecipitation with anti-CTCF antibodies and the immunoblotting for phospho-RxxS/T was performed. For quantification, the intensity of phospho-CTCF versus total CTCF signal in control samples is shown (right panel). **B.** T47D cells were transiently transfected with wild-type (WT) and T374E/S402E (phospho-mimetic) CTCF flag-tagged constructs. Subsequently, cells were lysed, and the chromatin fraction was isolated to display the presence of flag-CTCF bound to the chromatin fraction. **C.** Cell growth assays performed in T47D cells expressing both wild-type (WT) and a phospho-mimetic variant of CTCF T374E/S402E. **D.** The levels of T1079/T1041p LATS1+2 signal in both shEmpty (control) and shLATS1 cells is shown. The noticeable decrease in the phospho T1079/T1041 signal and LATS1 (Figure 5C) suggests that the predominant portion of detected p-LATS corresponds to LATS1 in the 3D model. GAPDH is used as loading control.

**EV6.**
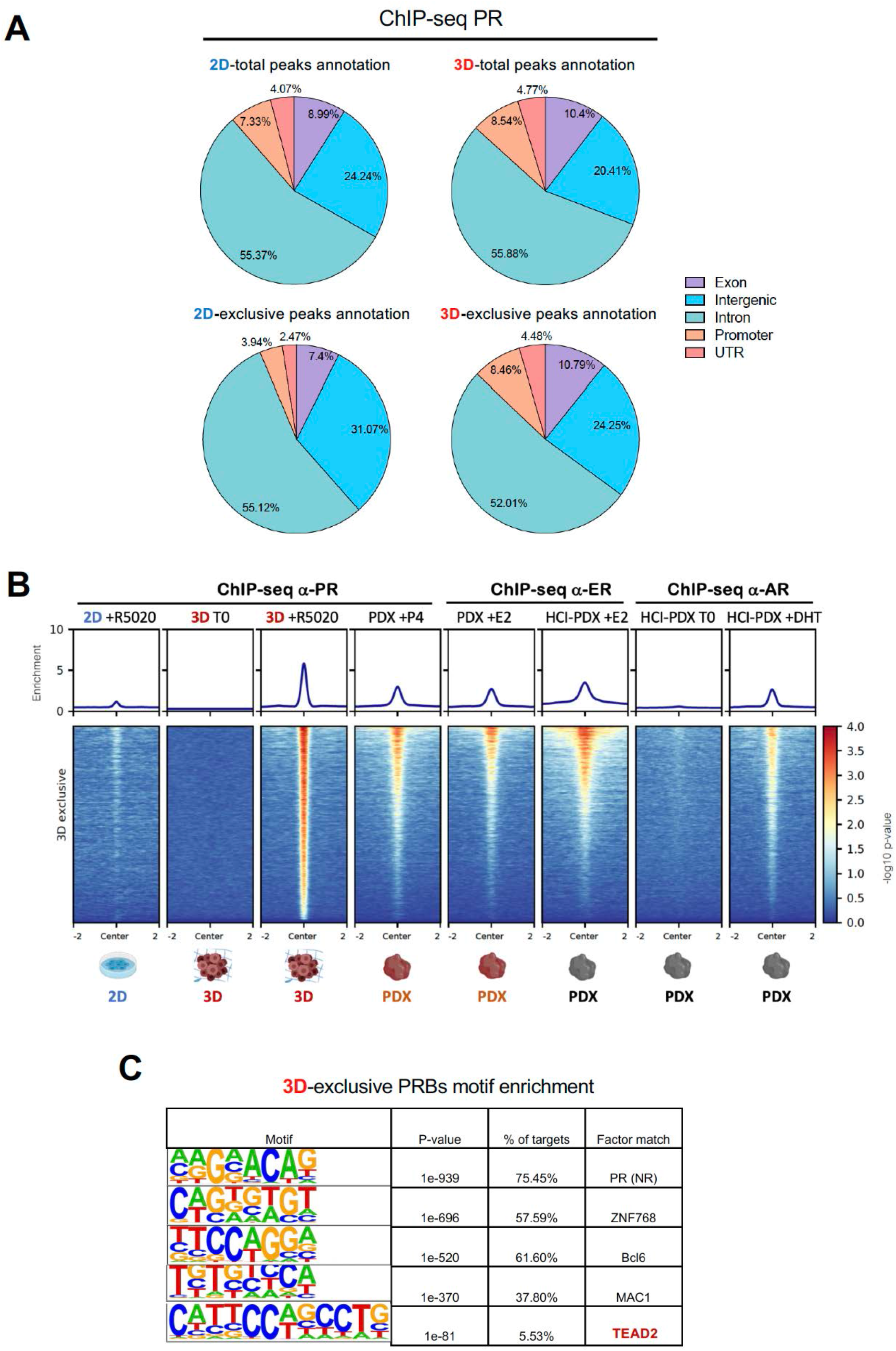
Hormone-induce PR binding in 2D and 3D grown T47D cells. Venn diagram of the 2D and 3D-exclusive PR binding sites (PRbs) from two replicates of ChIP-seq performed in 2D and 3D T47D cells in the presence and in the absence of 10 nM R5020 for 30 min. **B.** Progesterone (PR), estrogen (ER) and androgen receptor (AR) binding in the presence of their respective ligands in cells grown in monolayer (2D), in spheroids (3D) or in two patient-derived xenografts (PDXs). Nuclear receptor binding in the tumor is recapitulated exclusively by 3D cells. **C.** HOMER de novo motif enrichment analysis for 3D-exclusive PR peaks.

**EV7.**
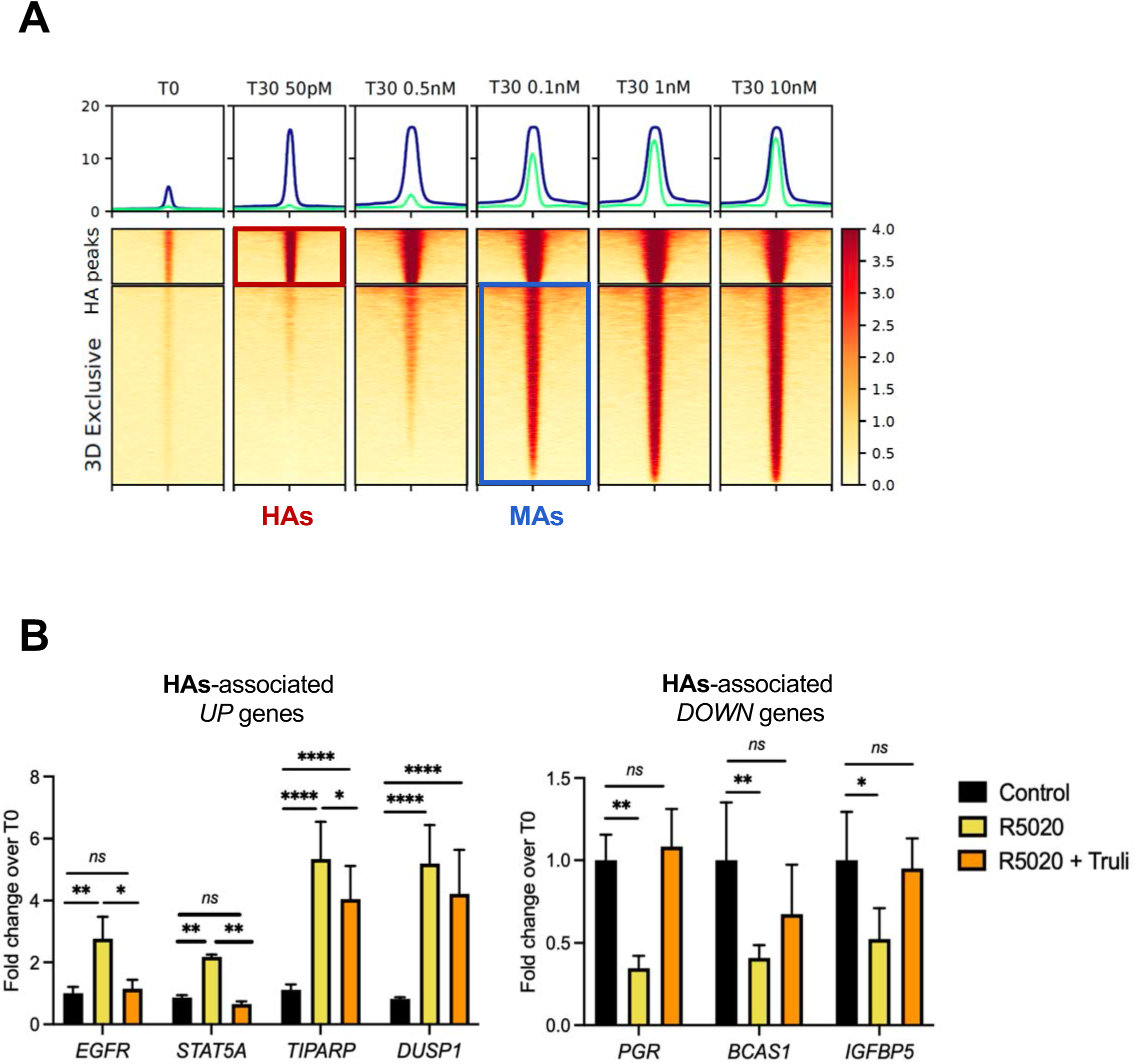
Hormone-dependent CTCF recruitment to High accessible PR binding sites (HAs) requires LATS1 activity. **A.** Heatmaps of PR ChIP-seq signal obtained at different concentrations of R5020 and corresponding to HAs and 3D-exclusive PR binding sites are shown. **B.** Cells grown in 3D conditions and treated or not with R5020 and TRULI as indicated, were submitted to gene activity assays. Four up and three down-HAs-associated genes were tested. Results are represented as mean and SD from two experiments performed in duplicate. The P value was calculated using the Student’s t-test.

**EV8.**
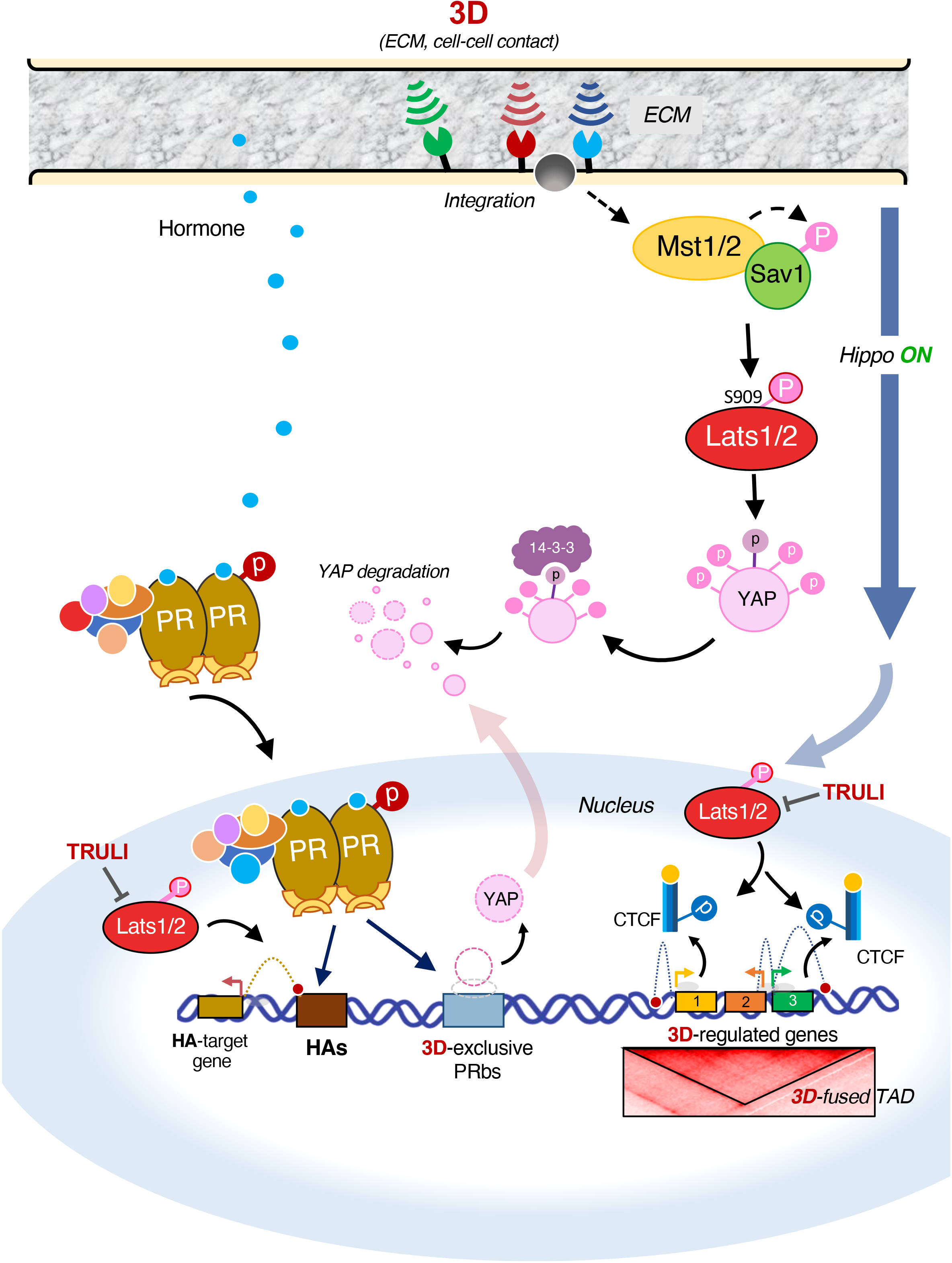
The Hippo kinase LATS1 controls CTCF binding and YAP nuclear availability in three-dimensionally grown breast cancer cells. T47D cells grown as spheroids are exposed to cell-cell contacts and to ECM. 3D-activated signals such as the Hippo pathway LATS kinase, impact on the cell nucleus in at least two ways: 1) the LATS1 kinase phosphorylates YAP promoting its cytoplasmic retention/degradation and 2) LATS1 also phosphorylates CTCF inducing its displacement from chromatin. The absence of these two proteins in the 3D nucleus determines the activity of a subset of genes specifically regulated in 3D condition and in turn, enhance the hormonal response.

## Appendix

**Figure S1.**
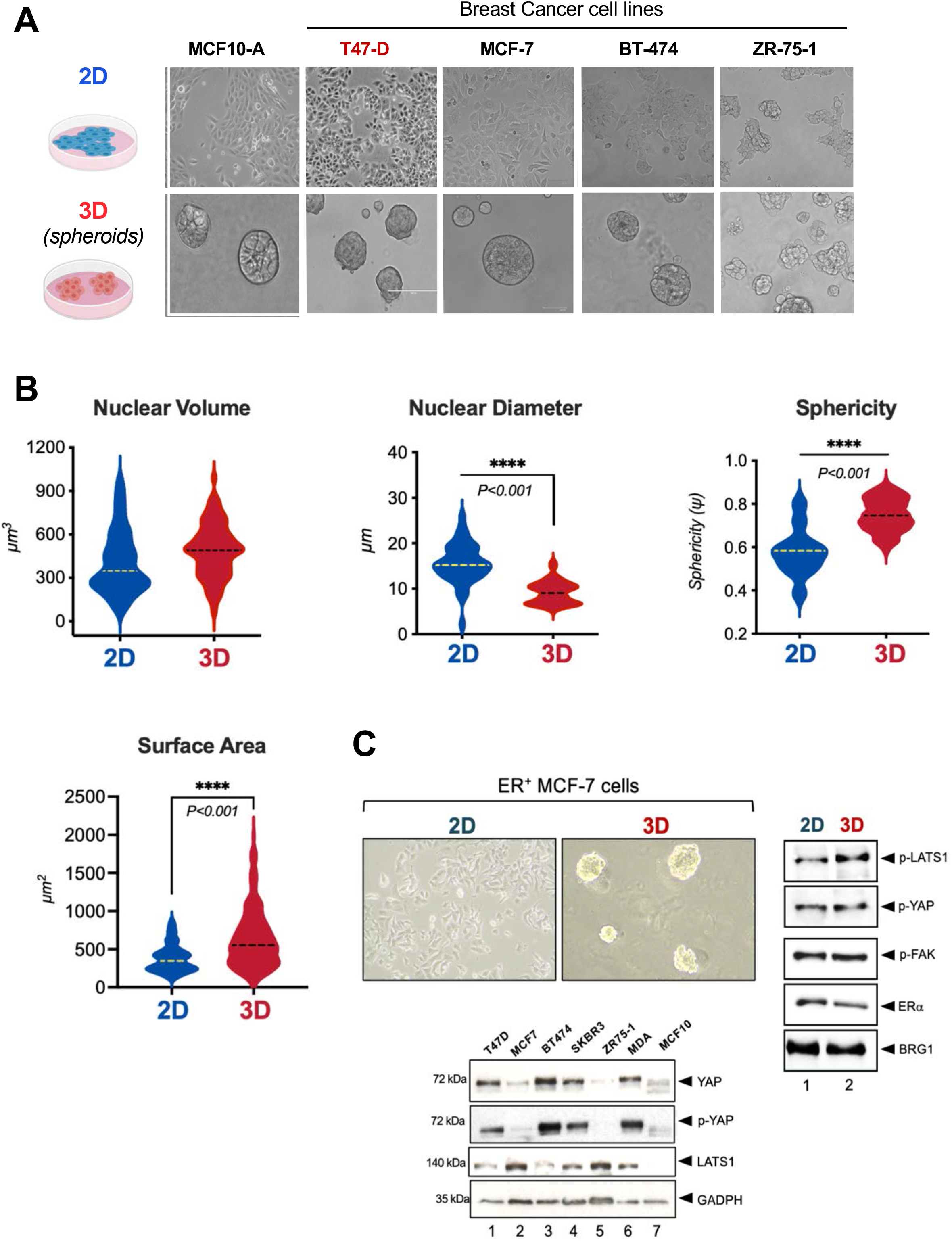
Breast cell lines grown in 3D culture embedded in Matrigel. Comparison of different breast cell lines grown on the plastic dish (2D) or as 3D spheroids in Matrigel. All cell lines in 3D can create defined spheres except for ZR75 which forms grape-like structures. Scale bar: 50 µm. **B.** *Physical properties of 3D cells.* The nuclei of the 3D cells measured with Imaris/ImageJ showed a larger volume accompanied by an increase of sphericity and larger surface area compared to 2D nuclei. **C.** MCF-7 cells were grown in 2D and 3D and the levels of p-LATS, p-YAP, p-FAK, ER and BRG1 was determined (right panel). High heterogeneity in YAP, p-YAP and LATS was found between different breast tumoral (lanes 1-6) and non tumoral (lane 7) cell lines.

**Figure S2.**
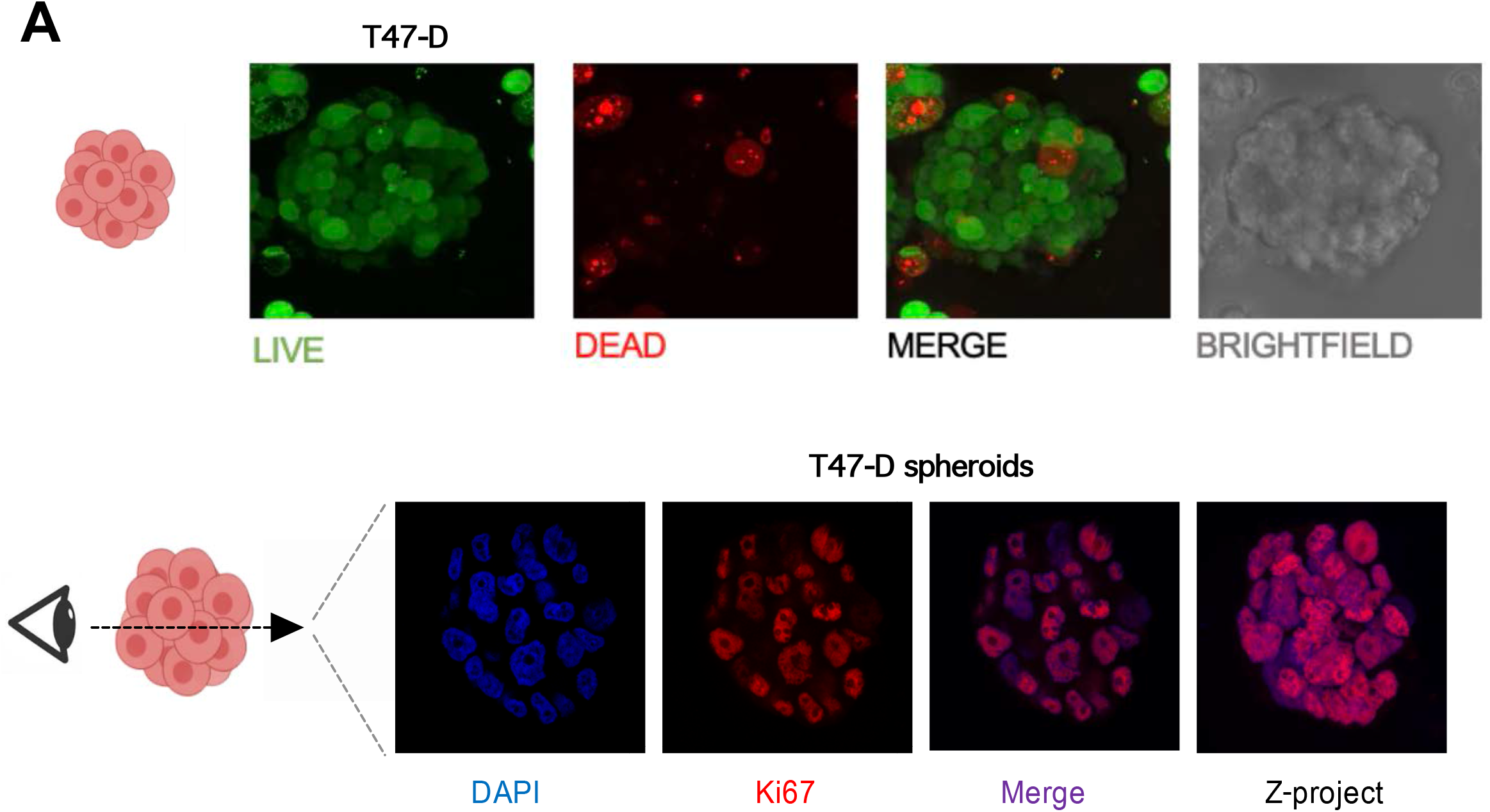
Cell proliferation in T47D spheroids. **A.** 3D T47D spheroid stained for live/dead cells after 10 days in culture. Apoptotic cores are rather small, sporadic and not predominant in the core of the spheroid. Bottom: Slice of a 10-day spheroid that was stained for the proliferative marker Ki-67 and overlapped with DAPI. Images were acquired on a Leica SP8-STED confocal laser-scanning microscope using the software Leica. Application Suite X with filters 488 (green) and 546 (red).

**Figure S3.**
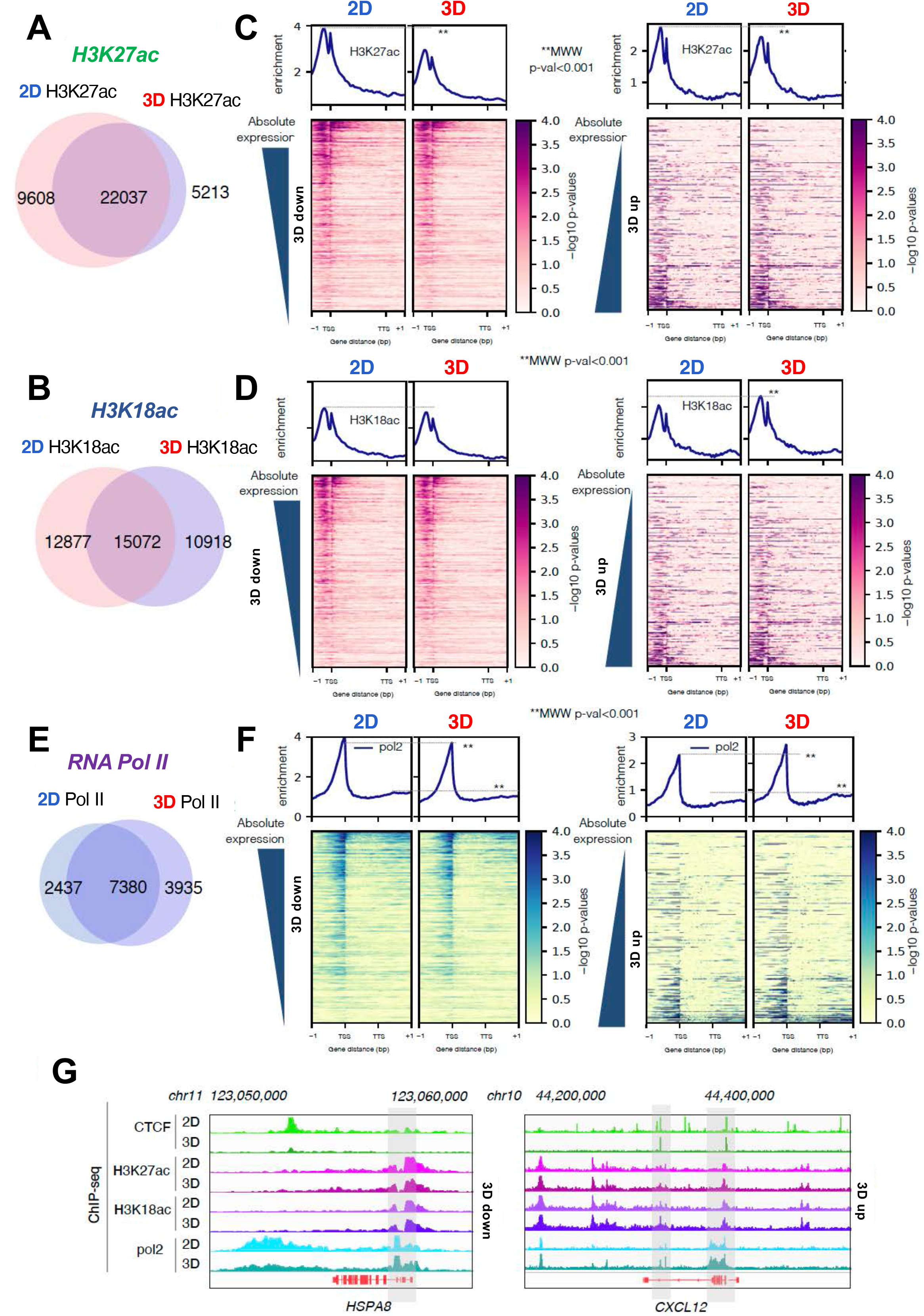
Epigenetic marks and RNApol Il distribution in 2D and 3D grown cells. Venn diagrams of H3K27ac (**A**), H3K18ac (**B**) and RNApol II (**E**) assayed in 2D and 3D grown cells. The H3K27ac (**C**), H3K18ac (**D**) and RNApol II (**F**) profiles in genes repressed or activated in 3D cells (left and right panels, respectively) is shown. **G.** Genome browser view of ChIP-seq data of *HSPA8* and *CXCL12,* two representative genes of each condition.

**Figure S4.**
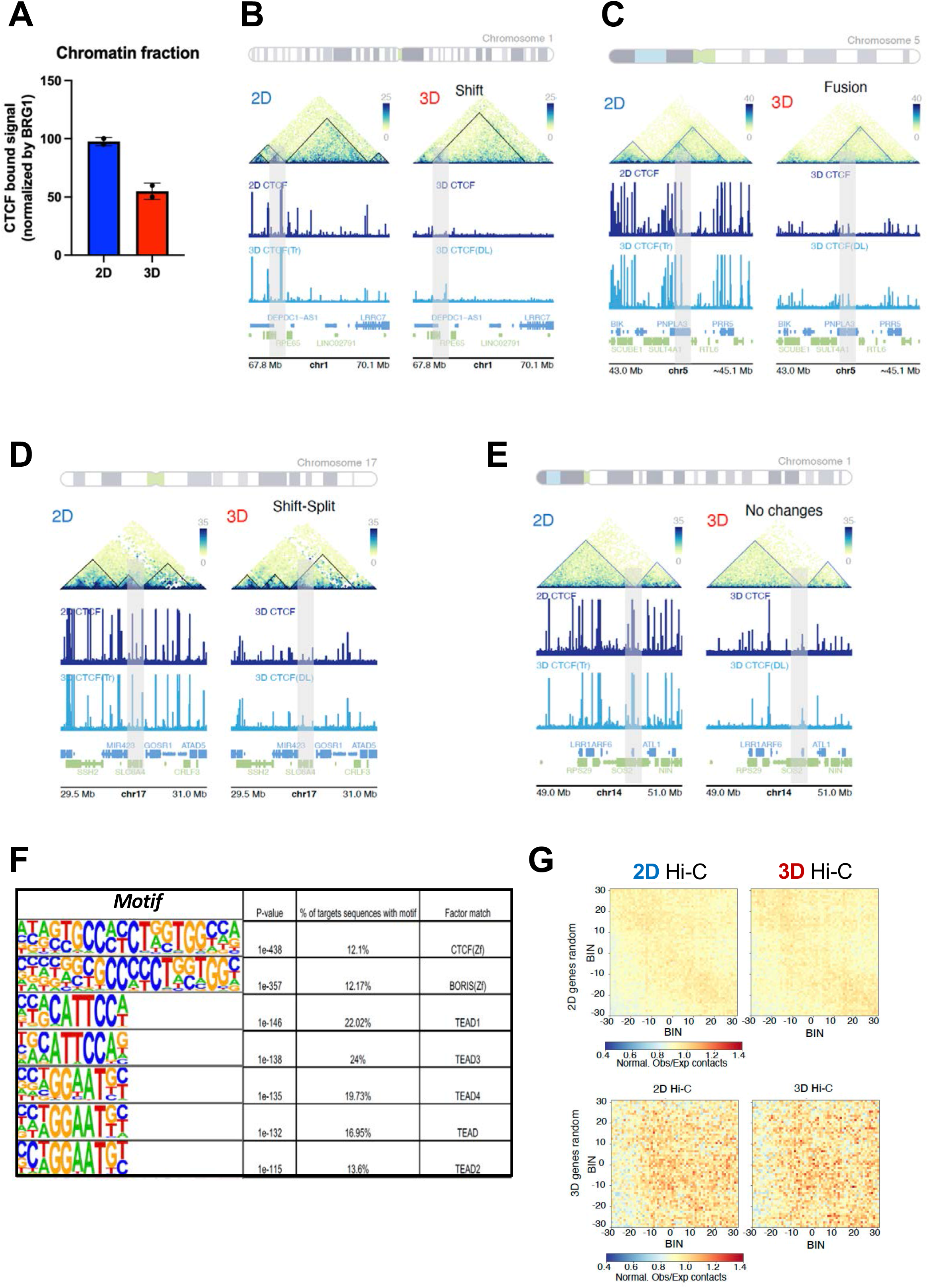
Loss of CTCF binding detected in 3D cells is not esclusively found at TAD fusions. **A**. T47D cells grown in 2D and 3D conditions were subjected to cell fractionation experiments (Mendez and Stillman, 2000). The chromatin fractions obtained in both conditions were run in polyacrylamide gels and western blot for CTCF was performed. The CTCF bound to chromatin was quantified using ImageJ software and the signal was corrected by the presence of BRG1. **B.** Loss of CTCF binding overlaps with: i) 3D-exclusive TAD split and DEG (green), ii) TAD fusions (blue), iii) unchanged TAD borders (gray) and iv) TAD fusions without any change in CTCF (purple). **C.** Loss of CTCF overlaps with TAD shifts and DEG (yellow) as well as with unchanged TAD borders (gray). **D.** Loss of CTCF binding without any changes in TAD border is shown (gray). **E.** Motif analysis of the ATAC-seq peaks: HOMER motif analysis identifies significant enrichment of the CTCF, CTCFL and TEAD motifs in ATAC-seq peaks with decreased signal in 3D. **F.** Hi-C explorer aggregate plots. Long-distance interactions among 3D and 2D random genes. The genomic coordinates of the random genes are centered between half the number of bins and the other half number of bins. Plotted are the submatrices of the aggregated contact frequency for 20 bins (1.5 kb bin size, 35 kb in total) in both upstream and downstream directions. Color bar scale with increasing red shades of color stands for higher contact frequency.

**Figure S5.**
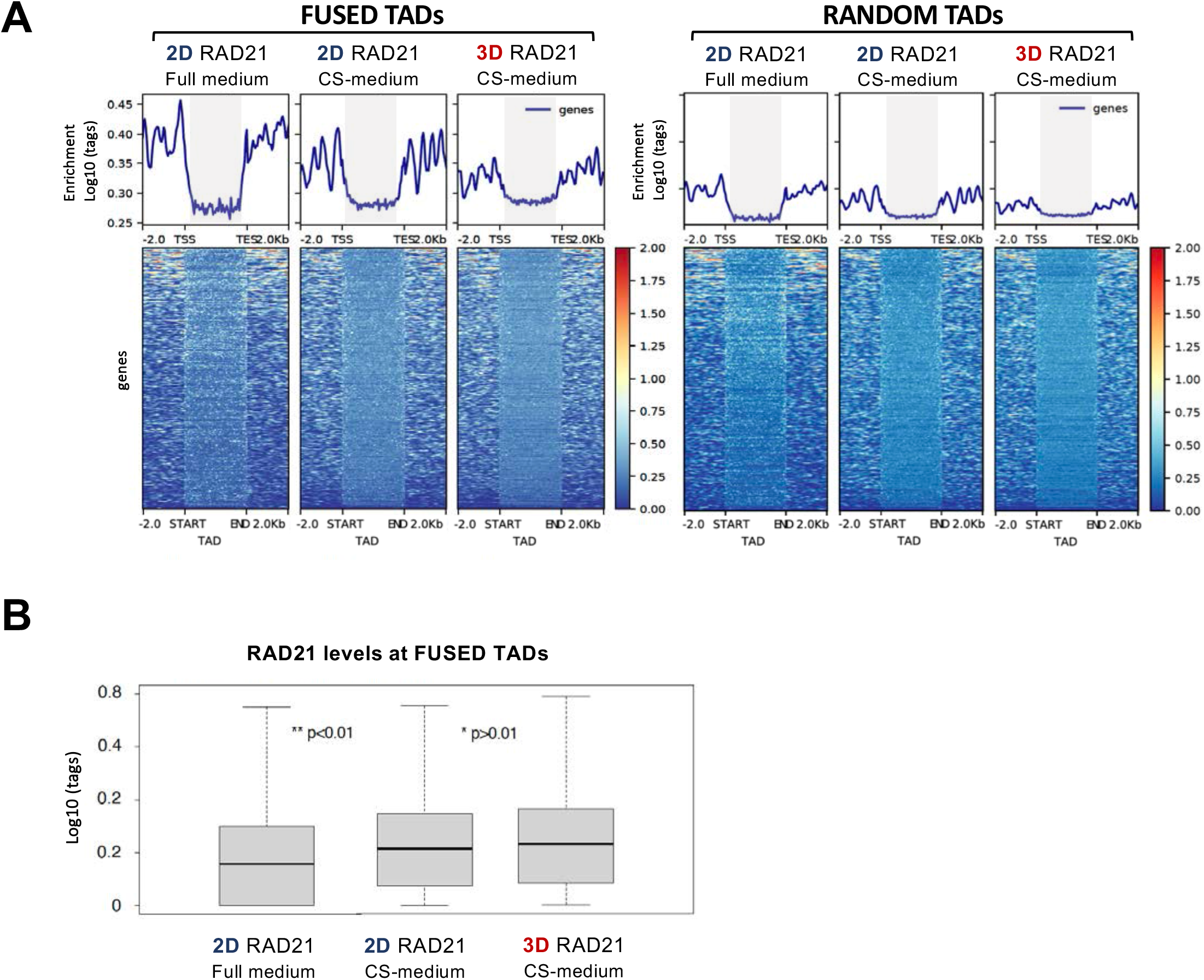
RAD21 distribution in 2D and 3D grown T47D cells. **A**. Heat maps of the cohesin component RAD21 ChIP-seq data around 3D-fused (left panel) and random Topological Associated Domains (TADs) (right panel). **B.** Box plots of the Log10 (tags) around each category. 2D: monolayer, 3D: spheroids, Full medium, CS: medium containing 10% charcolized serum without phenol red.

**Figure S6.**
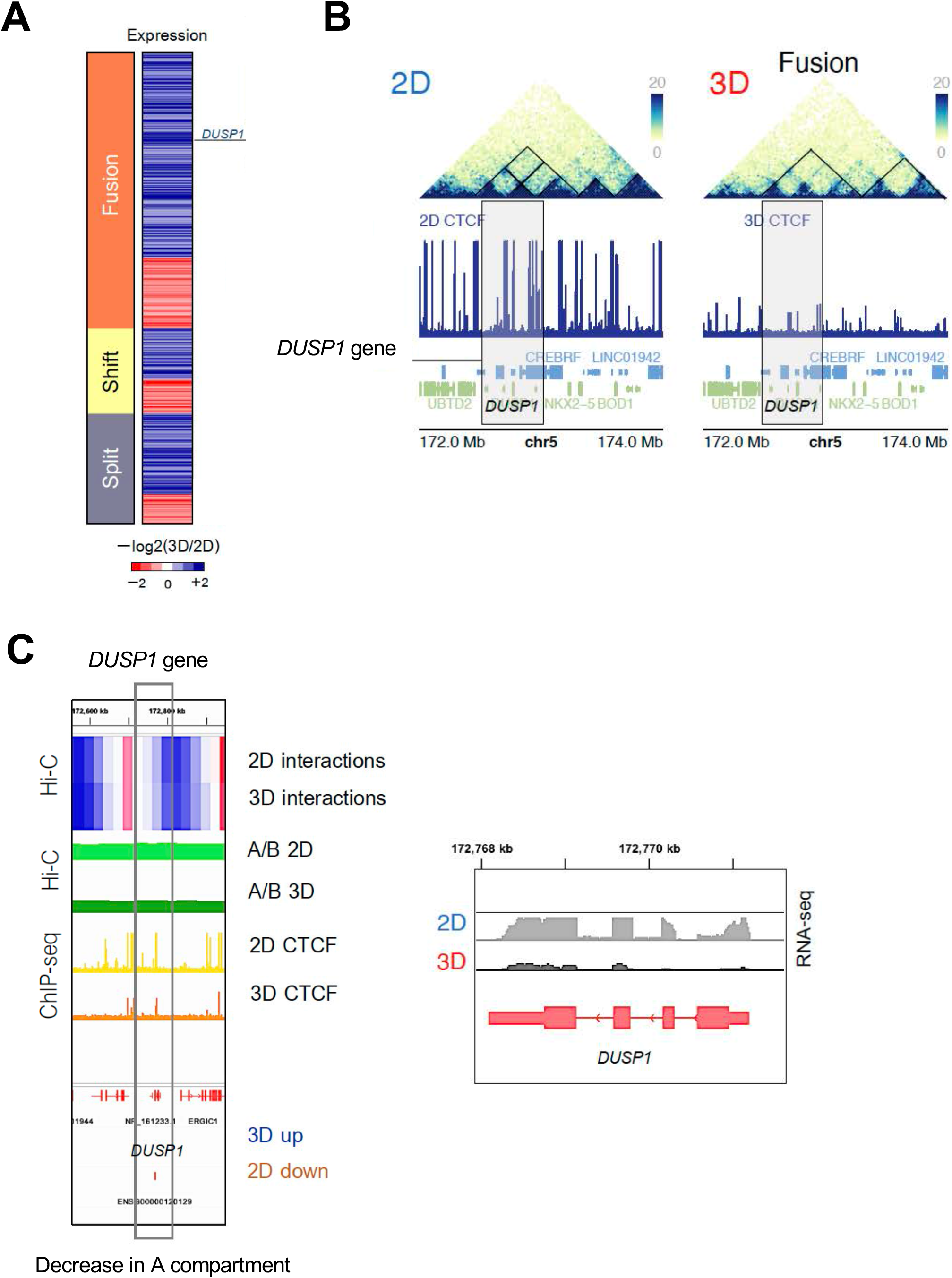
Effect of LATS1 inhibition on cell proliferation of 3D T47D cells. Cells grown in 2D and 3D conditions and treated or not with TRULI as indicated, were assayed for cell proliferation (**A**) and sphere size measurement after 10 days of culture (**B**).

**Figure S7.**
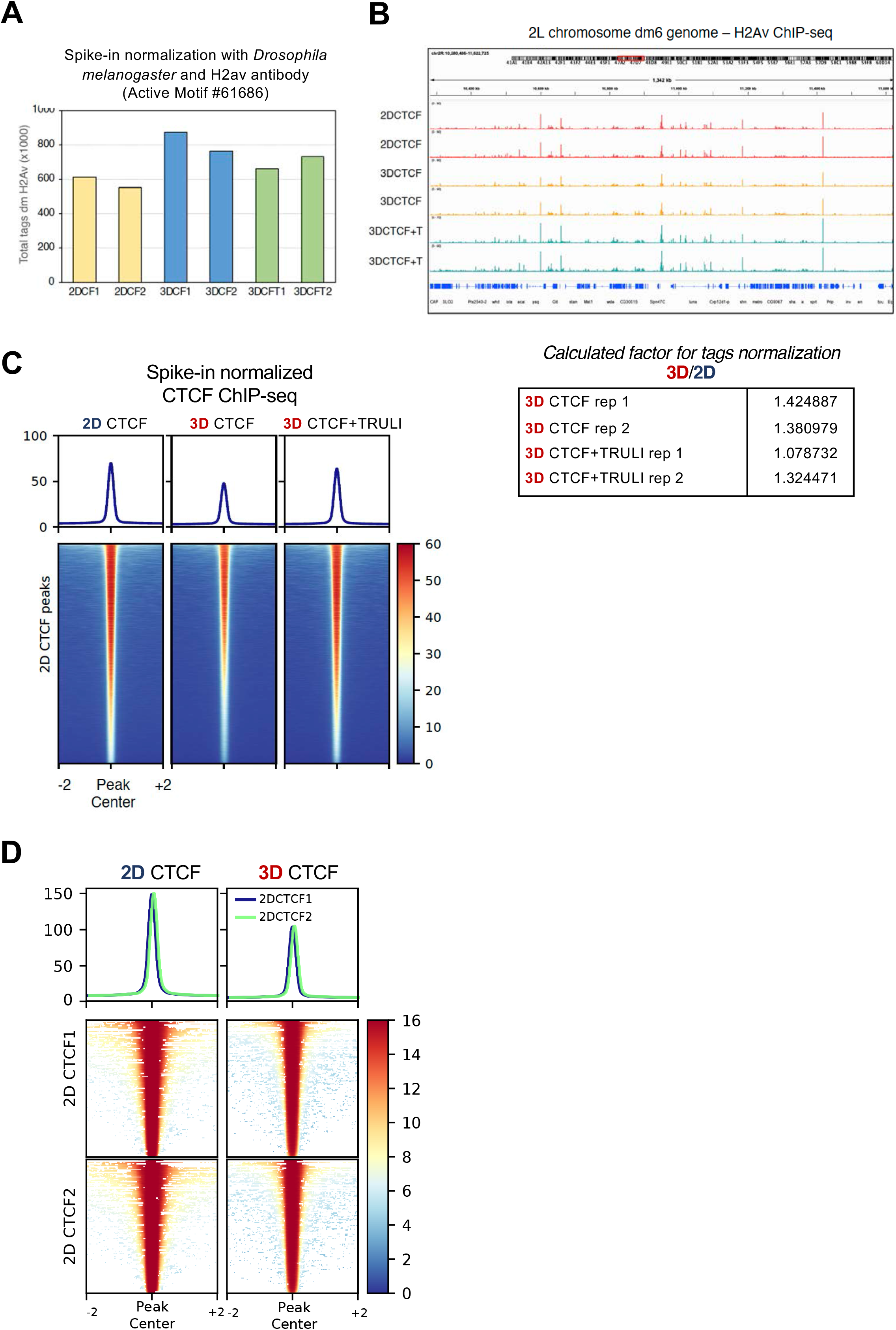
CTCF binding in 2D and 3D grown T47D cells using spike-in controls. **A.** Spike in normalization using Drosophila melanogaster chromatin and anti H2Av antibody (Active Motif) for CTCF ChIP-seq performed under mild conditions in 2D (yellow bars), 3D (blue bars) and 3D+TRULI (green bars) T47D cells. **B.** Snapshot of the genome browser showing the profile of H2Av ChIP-seq around the Drosophila 2L chromosome. **C.** Heatmaps of the CTCF ChIP-seq signal performed in 2D, 3D and 3D+TRULI cells and normalized using Drosophila melanogaster spike-in controls. **D.** *Heatmap of merged CFCT binding profiles*. CTCF ChIP-seq profiles from 2D and 3D grown cells have been merged to achieve an average value across replicates. BigWigMerge (UCSC genome tools) has been employed to merge all the bigwig profiles of all three replicates and generate one unique bigwig file. deepTool was implemented to compare newly generated merged bigwig to set of two replicates of 2D CTCF peaks. Significance of decrease has been measured by two-way ANOVA test.

**Figure S8.**
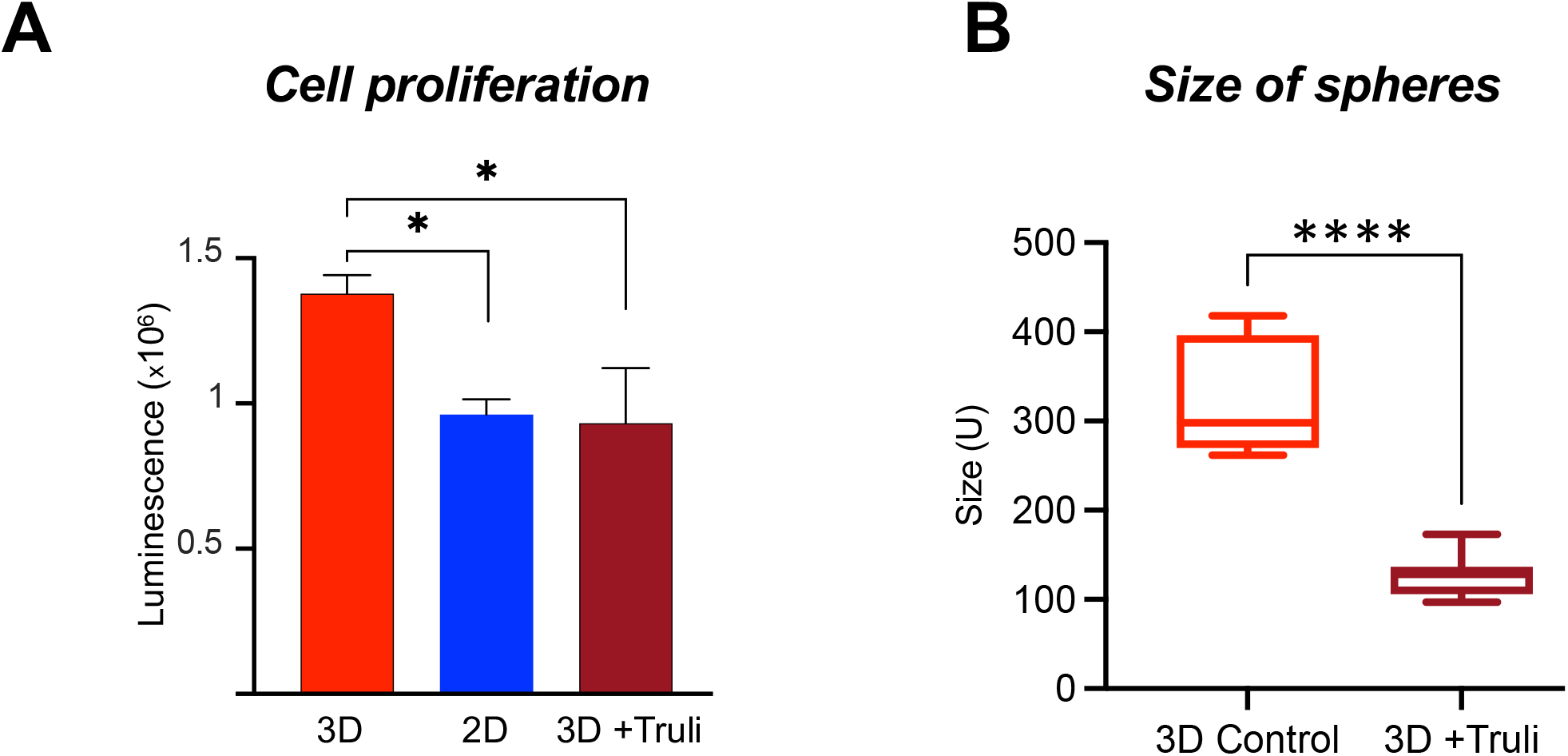
Activation of Progesterone Receptor in 2D and 3D T47D cells. Activation of Progesterone Receptor in 2D and 3D T47D cells. **A.** phosphoPR S294, total PR, ERa and FOXA1 levels were monitored in 2D and 3D cells treated or not with 10 nM R5020 for 30 min. **B.** PR activation between 2D and 3D conditions were assayed (left). The percentage of T47D cells that turned out to be responsive to hormone in both culture systems is shown. **C.** Immunostaining of 3D and 2D cells in the presence and in the absence of 10 nM of R5020 for 30 min, DAPI (blue), phospho-PR S294 (green), e-cadherin (red), and merge of all channels. **D.** PR and ERa expression in T47D cells grown under 2D and 3D conditions. Distribution of ERa, PR and b-catenin were assayed by immunofluorescence (IF) in 2D and 3D conditions. Scale bar: 20µm (left panels). Levels of PR and ERa between conditions was evaluated by western blot using specific antibodies (right panel).

**Figure S9.**
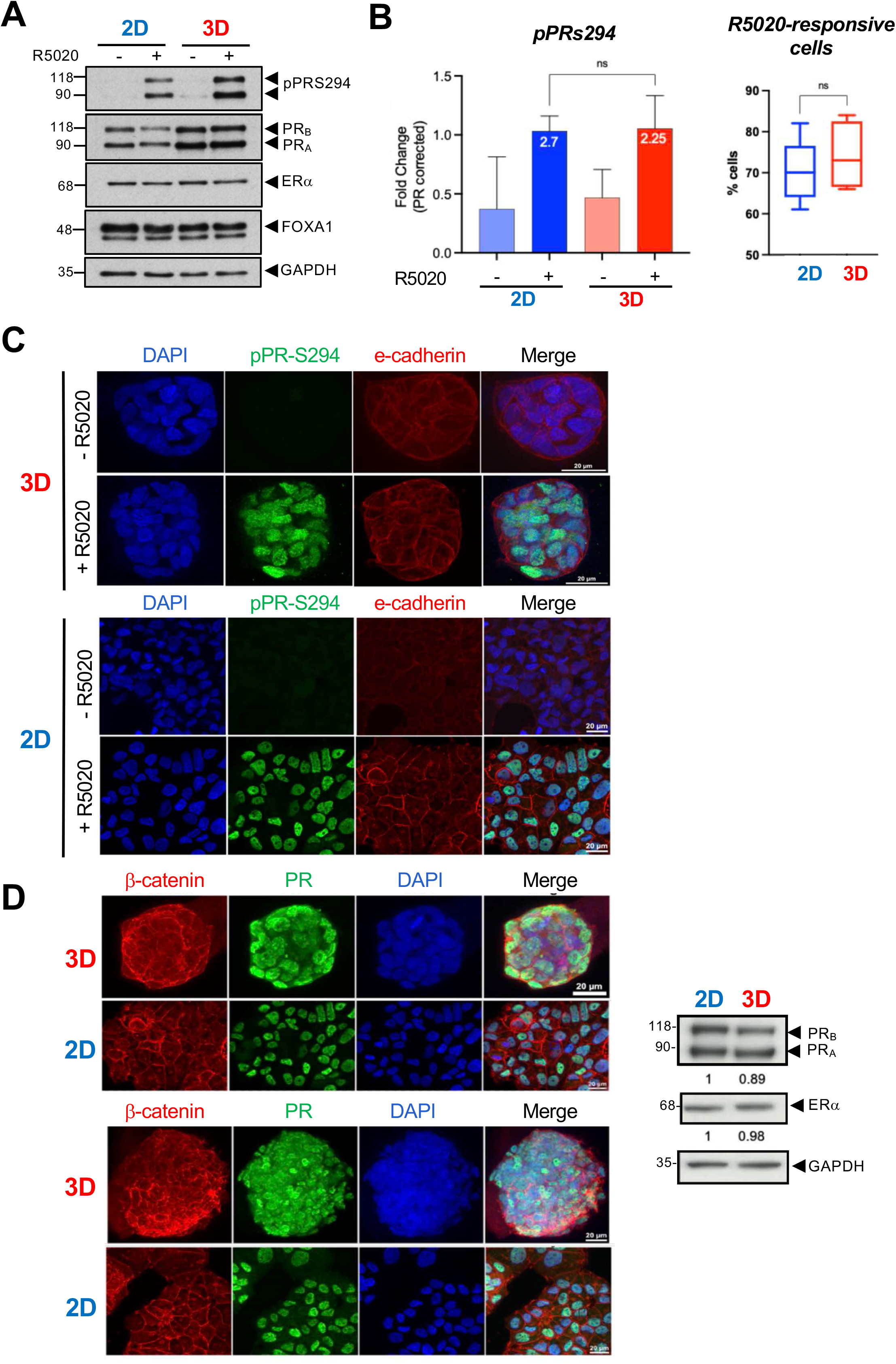
Enriched GO terms in 3D T47D cells treated with R5020. Using DEA results (Figure 6A), GO analysis was performed on 3D differentially expressed genes. The top terms are shown. The main enriched terms for up-regulated genes are associated with *cell structure, neurogenesis* and *neuron development*, while for down-regulated genes categories are related to *cell membrane components* and *cell contraction*.

**Figure S10.**
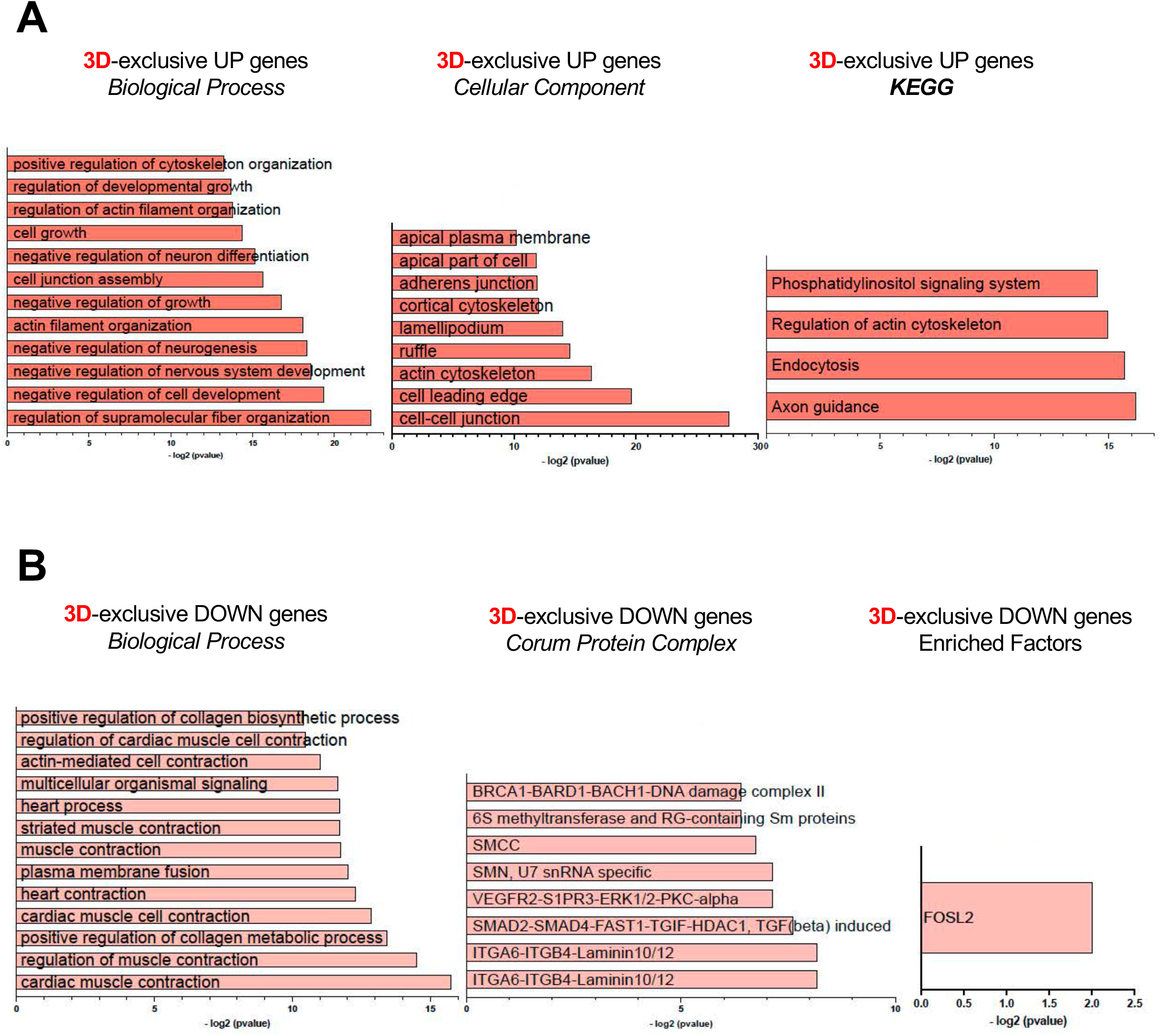
Hormone-induce PR binding in 2D and 3D grown T47D cells. **A.** Venn diagram of the 2D and 3D-exclusive PR binding sites (PRbs) from two replicates of ChIP-seq performed in 2D and 3D T47D cells in the presence and in the absence of 10 nM R5020 for 30 min. **B.** HOMER *de novo* motif enrichment analysis for 3D-exclusive PR peaks.

**Figure S11.**
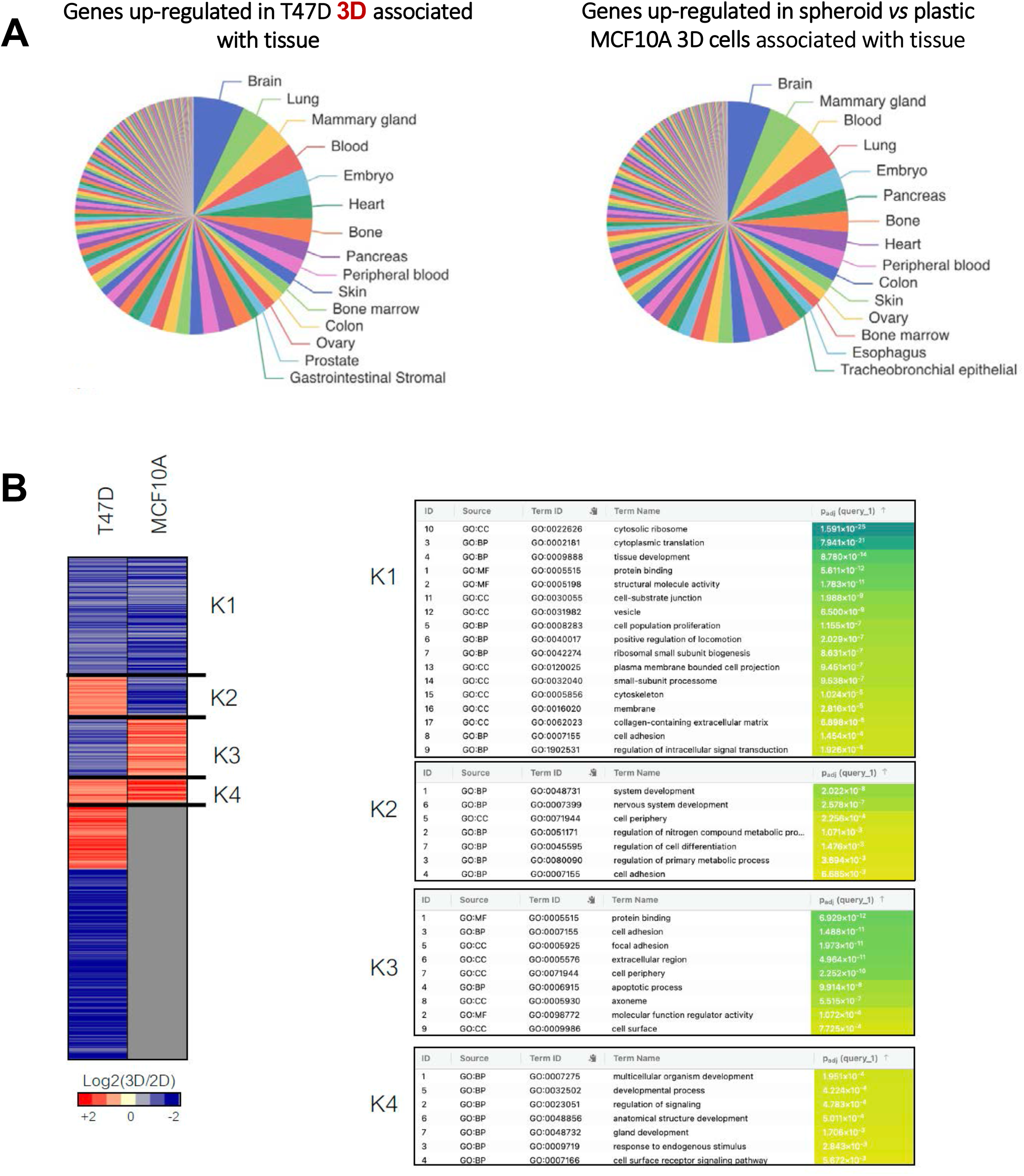
Gene expression profiles of tumoral and non-tumoral mammary cells when shifted to a 3D environment. **A.** Tissue association of up-regulated genes in T47D and MCF10A cells grown in 2D and 3D conditions. **B.** Gene clustering performed in T47D and MCF10A cells grown in 2D and 3D conditions is shown. *K2* and *K3* represent genes that are differentially expressed in tumoral *vs* non-tumoral model system once grown as spheres.

**Figure S12.**
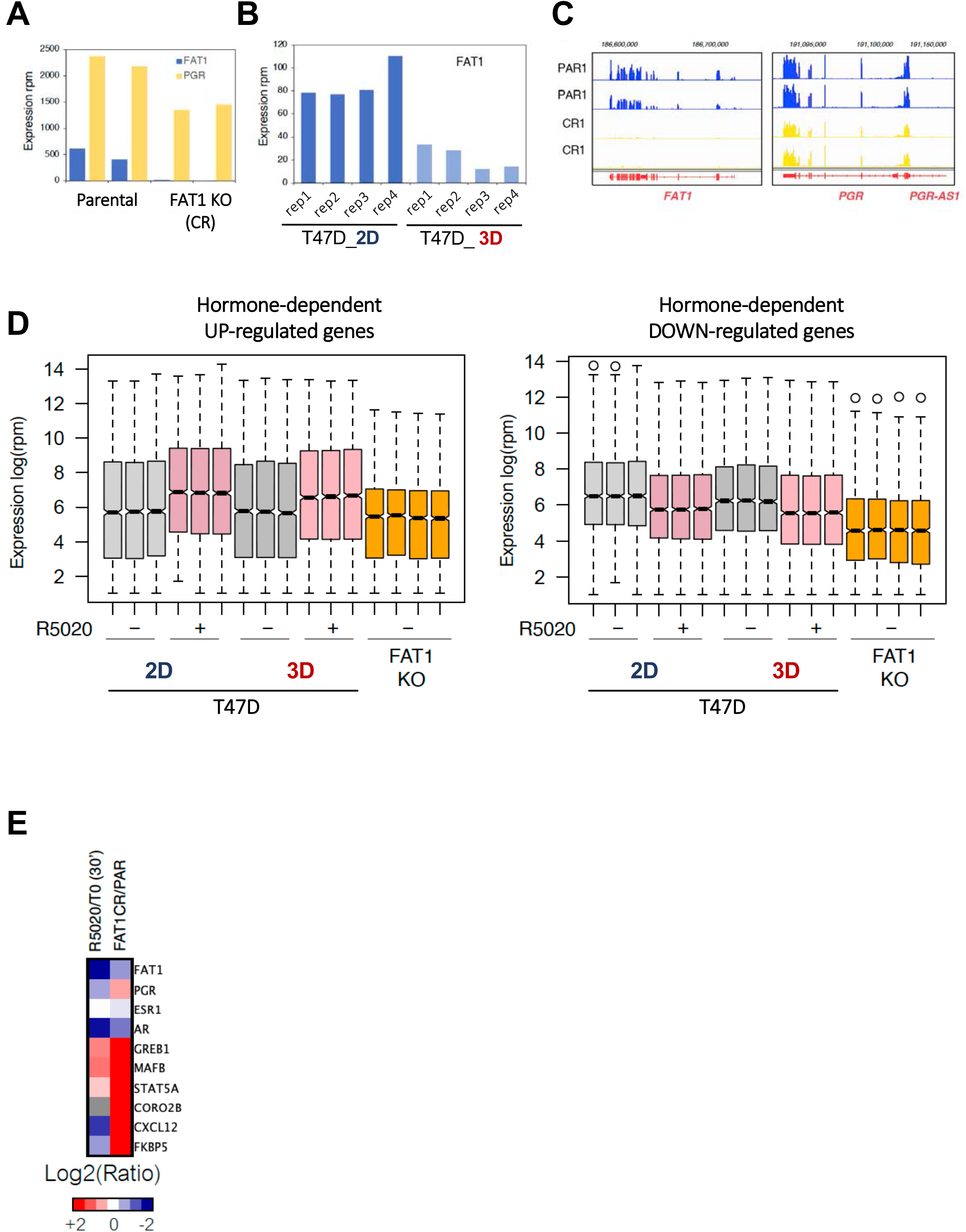
Connection between FAT1 KO and 3D-grown T47D cells. **A.** *FAT1* and *PGR* levels obtained from Parental and FAT1KO cells is shown. RNA-seq data was obtained from Li et al., Cancer Cell 34, 893-905 e8 (2018) and analyzed as described in extended bioinformatics methods section. **B.** FAT1 mRNA levels in 2D and 3D T47D cells. **C.** Snapshot of the genome browser showing the FAT1 and PGR expression in Parental and FAT1KO cells. **D.** Hormone-dependent expression in 2D, 3D T47D and FAT1KO cells. Genes up- and down-regulated in the presence of R5020 are shown (left and right panels, respectively). The trend presented for the same genes in FAT1KO cells is also shown. **E.** List of genes differentially regulated by R5020 and FAT1.

